# Centromeric origin of megabase-scale *Arabidopsis* interstitial telomeric repeat domains

**DOI:** 10.64898/2026.05.10.723265

**Authors:** Ivona Biocanin, Alejandro Edera, Clotilde Garrido, Adrien Vidal, Delphine Dardalhon Cuménal, Robin Burns, Katrin Fritschi, Jiří Fajkus, Detlef Weigel, Fernando A. Rabanal, Clara Bourbousse, Ian R. Henderson, Zhou Xu, Leandro Quadrana, Fredy Barneche

## Abstract

The mechanisms underlying the emergence of new DNA repeat classes remain poorly understood. One such example is the formation of interstitial telomeric repeat (ITR) domains commonly viewed as telomere-fossil byproducts of chromosome rearrangements. Here, we report that in *Arabidopsis thaliana*, ITRs arise from centromeres as centromeric-telomeric mosaic satellites that expand into higher-order structures via *in situ* amplification. Long-read sequencing of mutation accumulation lines and forward simulations of mutational processes confirmed the propensity of centromeres to form telomeric motifs. During establishment, ITRs adopt an atypical chromatin state that excludes CENH3 incorporation, thereby enabling escape from the functional constraints of centromere identity. Accordingly, a pangenomic survey of natural accessions showed that ITRs vary abruptly and independently of telomere length, notably through transposon-associated block duplications. These findings identify centromeres as major sources of telomeric sequences and an original route for the emergence of new DNA satellite types.

## Introduction

The advent of long-read sequencing technologies has enabled comprehensive profiling of telomere-to-telomere (T2T) genome sequences to cover highly repetitive chromosome regions partially or entirely missing from assemblies produced by Sanger sequencing or Next-Generation Sequencing (NGS). These developments recently allowed determining the entire sequence complement and exploring the diversification paths of megabase-long centromeres in mammalian and plant species^1,2^, or the Nucleolus Organizer Regions (NORs), composed of *45S* ribosomal DNA arrays in *Arabidopsis thaliana*^3^. However, the extent of interstitial telomeric repeat (or Sequence; ITR/ITS) domains in the genome across eukaryotes and their evolutionary history remain substantially unexplored, despite being identified decades ago in vertebrate nuclei^4^ and detected in multiple species using *in situ* hybridization of telomeric DNA sequences^5–7^. Highly repetitive ITR elements carrying a few to hundreds of telomeric repeat (TR) sequences make up as much as 5% of Chinese hamster nuclear DNA^8^ and likely represent more than 0.5 Mbp in the *A. thaliana* genome^9^. ITRs accumulate during carcinogenesis^5,6^ and can constitute hotspots for DNA damage and chromosome breakage^10^. Hence, unlike the chromosome-end-protective role of telomeres^11,12^, ITRs are associated with genome instability, and their function is poorly understood.

In *A. thaliana*, ITRs can form conspicuous chromatin subnuclear domains tethering Polycomb Repressive Complex 2 and impact genome topology^9^. In other species as well, ITRs may not constitute inert telomere byproducts but influence epigenome regulation by acting as recruitment substrates or sequestration sites for gene-regulatory proteins and chromatin domains^5,13^. Like their function, the origins of ITRs have long remained uncertain. Their common pericentromeric or sub-telomeric localization led to the prevailing view of ITRs as remnants of telomeric material internalized through chromosome rearrangements such as end-to-end fusions or Robertsonian translocations^5,14^, possibly acting as a substrate for the formation of a new telomere or an acrocentric chromosome^4,15^. Yet, studies on numerous species often failed to identify ITR footprints at chromosome fusion sites^7,16^.

Here, we address this paradox by exploring ITR architectural variations in *A. thaliana* using a complete genome assembly, sequencing data from mutation-accumulation lines and over 800 natural accessions, uncovering their emergence from centromeric satellites and two complementary mechanisms that underlie their rapid evolutionary dynamics.

## Results

### Telomeric sequences at centromeres and ITRs

We conducted an exhaustive search of TR sequences in the *TAIR10* reference genome and four long-read-based assemblies of the *A. thaliana* reference accession Col-0 (Extended Data Fig. 1). Using all frames of the AAACCCT 7 bp-long consensus telomeric motif in both orientations, and omitting the non-assembled telomeres *TEL2-L* and *TEL4-L* that cap *NOR2* and *NOR4*, we identified about 140,000 TRs per genome; a minority of which (approximately 3,800) constitute terminal telomeres of 2.9 to 3.8 kb (Extended Data Fig. 1). To avoid potential telomere assembly biases and capture the missing lengths of *TEL2-L* and *TEL4-L*, we inspected PacBio HiFi sequencing reads from the Col-CC resequencing dataset (Supplementary Table 1) using *TeloReader*^17^ (Extended Data Fig. 1). In line with possible chromosome arm-specific telomere length (TL) setpoints^18^, the 10 telomeres display broad variations of average length, *TEL4-L* being the shortest (2.1±0.35 kb) and *TEL2-L* among the longest (3.5±0.9 kb) (Fig. 1a, Extended Data Fig. 1). Overall, telomeres account for ∼2.7% of the TR library while the vast majority correspond to interspersed elements (Extended Data Fig. 1).

**Fig. 1.**
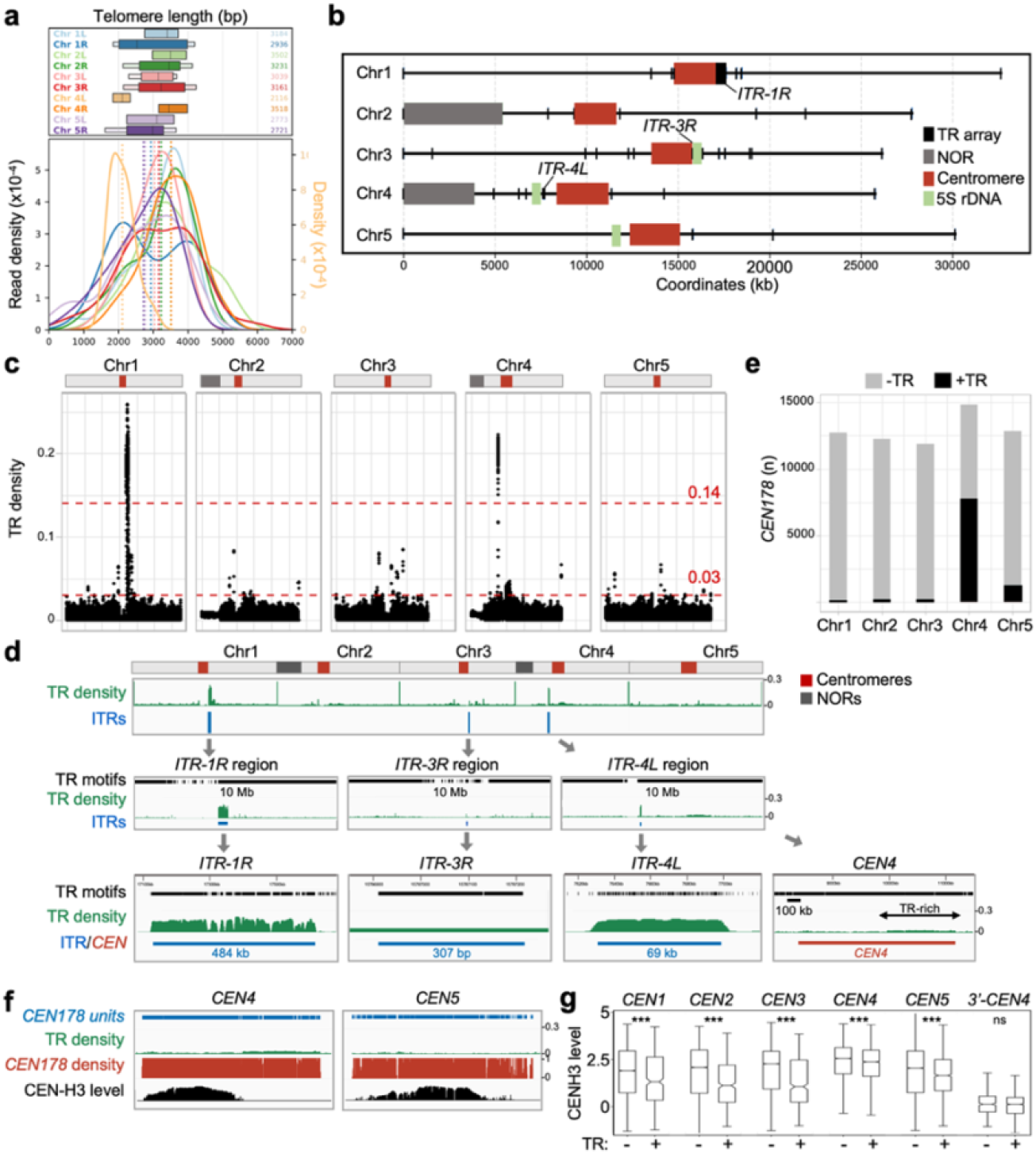
Telomeric motifs substantially populate ITRs and centromeric satellites in *A. thaliana*. **a**, Chromosome end-specific telomere length distributions determined using *Col-CC* long sequencing reads are represented as colored boxplots and densities. The right y-axis scale corresponds to chromosome *4L* telomere, and the left y-axis to all other chromosome extremities. The number of telomere-associated reads is indicated. **b**, Chromosome map of TR arrays (>20 bp) in *Col-CC*, allowing on average 1 SNP every 23 bp. **c**, TR density across *Col-CC* chromosomes using a 5-kb window and TR density thresholds distinguishing the major ITRs *ITR-1R* and *ITR-4L* as well as a TR-rich *CEN4* terminal domain. **d**, Overview of TR-rich domains in Col-0. In addition to typical ITRs (*ITR-1R, ITR-3R*, and *ITR-4L*), peripheral regions of the *CEN4* and *CEN5* centromeres exhibit a high TR density with, on average, one canonical telomeric motif per *CEN178* satellite variant. **e**, Number of *CEN178* satellites carrying (+) or not (-) canonical TRs per chromosome. Centromeric domains were defined as regions bearing clustered *CEN178* consensus sequences^2^, not limited to CENH3-occupied regions. **f**, ChIP-seq profile showing that CENH3 level drops at TR-containing *CEN4* and *CEN5* regions. **g**, CENH3 levels (IP/Input Log2 fold change) per *CEN178* satellite carrying (+) or not (-) telomeric motifs. ***, p-Value<10^-5^; ns, p-Value>10^-2^.

To identify clusters of interspersed TRs characteristic of ITRs, we first searched the Col-CC T2T genome for domains containing at least three consecutive TR motifs (see Methods). This approach identified short TR arrays in pericentromeric regions, with the longest one, on the right arm of chromosome 3, measuring ∼300 bp (*ITR-3R*; Fig. 1b, Extended Data Fig. 1). We also searched for heterogeneous ITRs in which consensus TRs are interrupted by degenerated or unrelated sequences by implementing *ITR-scan*, a pipeline that detects TR-rich domains within 5-kb genome windows regardless of TR consecutiveness (Extended Data Fig. 1). This identified two outstandingly long ITRs with significant TR density. While *ITR-1R* (484 kb, 9987 TRs) lies immediately adjacent to the right border of centromere 1 (*CEN1*), *ITR-4L* (69 kb, 1933 TRs) is separated from a *5S* rDNA array and the left border of *CEN4* by transcriptionally active genes and transposable elements (TEs) (Fig. 1b-1d, Extended Data Fig. 1). *ITR-1R* and *ITR-4L* sequences are largely conserved across three independently generated T2T genome assemblies, showing inter-individual ITR stability (Extended Data Fig. 2).

Using a lower TR density cut-off, *ITR-scan* detected 38 other ITR candidate loci, altogether harboring approximately 5,000 TRs. The longest one spans a 1.14-Mb region of *CEN4*, in which most satellites carry a telomeric motif resulting from a single-nucleotide substitution from the *CEN178* consensus sequence (Fig. 1c-f, Extended Data Fig. 3). To assess whether TR presence influences centromere identity, we examined the genomic distribution of the CENH3 histone variant, a CENP-A homolog whose incorporation into chromatin defines centromere functionality in plants^19^. CENH3 levels are significantly lower at TR-containing *CEN178* satellites compared to other *CEN178* satellites (Fig. 1g) with a sharp attenuation across the TR-rich *CEN4* region (Fig. 1f, Extended Data Fig. 3). Together, these data indicate that point-mutation-driven emergence of TRs within *CEN178* satellites compromises local centromere function.

### Pericentromeric ITRs emerge in *cis*

To assess how the major ITRs form, we searched for their elemental repeat units. Consistent with ITRs being composed of monomeric units, analyses of inter-TR distances in *ITR-1R* and *ITR-4L* identified a sharp periodicity of 530 bp and 600 bp, respectively (Fig. 2a, Extended Data Fig. 3). At a finer scale, ITR periodic units carry ∼180-bp-long clusters of TRs, hereafter referred to as *Telo*-islands (Fig. 2b-c, Extended Data Fig. 3), which possibly correspond to the 180-bp and telomere polymorphic fragments identified in early analyses of restriction profiles and BAC clones of *A. thaliana* genomic DNA^20–22^. A template-based repeat analysis showed that spacer elements between *Telo*-islands contain one or more antisense *CEN178* centromeric satellites with 70-90% identity to the *CEN178* consensus sequence (Fig. 2c, Extended Data Fig. 3). Long ITRs are therefore composed of mosaic monomers, hereafter referred to as *CenTel* satellites, which combine telomeric and centromeric-related sequences in opposite orientations. *Telo*-islands typically abut a truncated *CEN178* satellite variant, regardless of the length of the *CenTel* unit (Fig. 2c, 2e), suggesting that TR clusters emerged from *CEN178* sequence drift, as observed at *CEN4* and *CEN5* with single interspersed telomeric motifs. We thus tested the hypothesis that *CenTel* units originate from their proximal centromere by exploiting the existence of private centromeric sequence polymorphism in each chromosome^2^. Sequence distance analyses of *CenTel* satellites relative to all chromosome-specific *CEN178* satellites showed that *ITR-1R*/*CEN1* and *ITR-4L*/*CEN4* satellite sequence pairs are significantly more similar than any inter-chromosomal *ITR*/*CEN* pair (Fig. 2f). Based on these observations, we conclude that each ITR originates from the neighboring centromere by local emergence of a new satellite type.

**Fig. 2.**
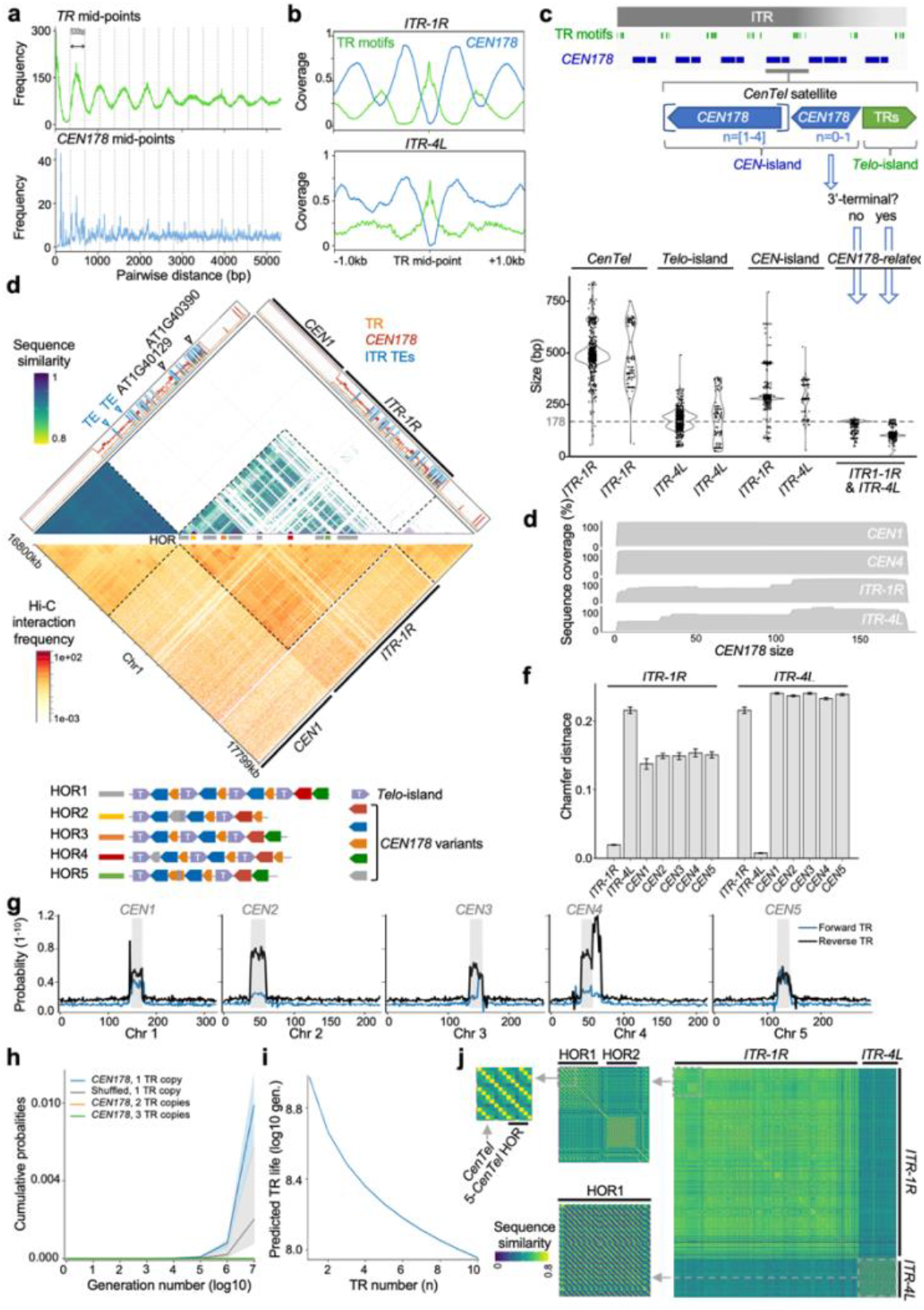
Long ITRs form as higher-order repeats of telomeric and centromeric mosaic satellites. **a**, Frequency of pairwise distances between mid-points of TRs and *CEN178* sequences in *ITR-1R*. **b**, Coverage of TR and *CEN178* satellite sequences within *ITR-1R* and *ITR-4L* relative to the TR sequence mid-points. **c**, Top panel, schematic representation of ITR *CenTel* monomer organization as clusters of canonical and related TR sequences (*Telo*-islands) interspersed between short arrays of 1-to-4 *CEN178* variants (*CEN*-islands), with TRs and *CEN178*-related sequences always oriented head to tail. Bottom panel, size distribution of *CenTel, Telo*-island, and interspersed *CEN178*-related sequences, showing that the 5’-end of terminal *CEN178* variants is truncated. **d**, *ITR-1R* internal sequence identity (top) and 3D organization (bottom). Top, TR and *CEN178*-like repeat density. Two of the TEs and TE-related genes embedded in *ITR-1R* are shown as triangles. Middle, Hi-C profiles at 1-kb resolution, showing that 2 TE copies (empty triangles) mark the border of insulated HORs. *ITR-1R* is insulated from *CEN1* and from heterochromatic domains (dashed lines). Bottom, depiction of major *ITR-1R* HORs and *CenTel* variations. *ITR-1R* is predominantly composed of 5- or 4-*CenTel* HORs interrupted by 2 or 3-*CenTel* HORs. **e**, Aggregated alignments of all *CEN178*-like sequences in *ITR-1R, ITR-4L* and their neighboring centromere to the consensus *CEN178* sequence^2^ showing the presence of truncated *CEN178* satellites in about one-third of the ITR *CEN178* variants. **f**, Chamfer distance between *CEN178* sequences within centromeres, *ITR-1R*, and *ITR-4L*. Error bars represent the variability estimated via bootstrap resampling (n=50). **g**, Chromosome-wide probability of TR emergence for all seven AAACCCT phase shifts in both orientations. **h**, Simulation of single (N=1) or multiple (N=2,3) TR appearance probability as a function of the number of generations. Lines show the median cumulative sum over time with shades indicating 30th and 70th percentiles. **i**, Expected life of TRs based on copy number. **j**, Distance matrix of *CenTel* units in *ITR-1R* and *ITR-4L. CenTel* ITR monomers are represented as dots and colored by sequence similarity, identifying yellow lines at HOR boundaries. *CenTel* coordinates were defined as *Telo*-island intervals.

To characterize *CenTel* satellite sequence dynamics, we performed PacBio HiFi genome sequencing for four mutation accumulation (MA) lines derived from the same founder plant propagated by self-inbreeding for 32 generations to determine point mutation (PM) rates and spectra genome-wide, including within the centromeres and ITRs (Extended Data Fig. 4). These lines are direct descendants of the cohort sequenced by Becker *et al*. (2011)^23^. This revealed that the *ITR-1R* PM rate is one order of magnitude higher than that of protein-coding genes (1.8×10^-8^ and 3.3×10^-9^ per bp per generation, respectively), reaching levels comparable to those of TEs (1.6×10^-8^), *CEN1* (2.8×10^-8^), or *CEN5* (2.3×10^-8^) (Extended Data Fig. 4). Using the centromere PM rate (μj=4.7×10^-8^) and excluding pre-existing TRs, we found that the probability of PM-driven TR formation is markedly higher within the five centromeres than at any other region of the genome (Fig. 2g, Extended Data Fig. 4). Moreover, the per-satellite probability of acquiring a single TR (1×10^-9^) is significantly higher than expected from scrambled *CEN178* sequences (permutation test; p-val=1e-4). In contrast, the probability of forming multiple TRs within the same *CEN178* satellite is negligible, suggesting that *Telo*-islands do not form solely via PMs (Fig. 2h).

Forward simulations of sequence evolution parametrized using these PM rates and spectra predicted substantial divergence (over 6%) from the *CEN178* consensus after 10 M generations (Extended Data Fig. 4) and long persistence of TRs (expected lifetime of ∼794 M generations), which decreases to ∼79 M generations for satellites carrying more than 10 TRs (Fig. 2i). With a high birth rate and long lifetimes, the model therefore predicts a net accumulation of *CEN178* satellites carrying TRs over generations, further supporting that TR-rich *CEN178* and *CenTel* satellites originate from centromeres.

At a broader scale, *CenTel* sequence-similarity analyses showed that *ITR-4L* consists of a single higher-order repeat (HOR) region while *ITR-1R* contains multiple and diversified HOR blocks (color blocks and chessboard patterns in Fig. 2d, 2j, Extended Data Fig. 5) primarily interrupted by TEs (white stripes in Fig. 2d). The presence of multiple HORs and *GYPSY*-type *ATHILA* retrotransposons is reminiscent of centromere organization^2^, yet *ITR-1R* is also enriched in *ENSPM* DNA transposons and *ATLANTYS* retrotransposons relative to their genome-wide occurrence (Extended Data Fig. 5). Chromosome conformation capture (Hi-C) showed that HOR boundaries generally define *ITR-1R* topological subdomains, and that ITRs appear insulated from the neighboring centromere (Fig. 2d, Extended Data Fig. 6). Together, these data indicate that *CenTel* emergence from centromeric satellites was followed by local spreading, sequence diversification, and spatial compartmentalization of the newly formed ITR.

### *CenTel* satellites evade centromere function

Given their mosaic nature, we envisaged that the chromatin status of *CenTel* satellites may relate to that of telomeres or centromeres. Like telomeres, *ITR-1R* and *ITR-4L* display a high propensity to form intramolecular or intermolecular G-quadruplexes, which may impair ITR stability (Fig. 3a, Extended Data Fig. 6), but unlike telomeres^11^, ChIP-seq profiling showed that Telomeric Repeat Binding (TRB) proteins 1, 2, and 3 are not enriched at ITRs.

CENH3 is also absent at *ITR-1R* and *ITR-4L* (Fig. 3a, Extended Data Fig. 6, Supplementary Table 2), consistent with its low enrichment at TR-containing centromeric domains (Fig. 1f, 1g). The two ITRs also lack hallmarks of euchromatin (H3K4me3) and of PRC2 activity (H3K27me3) but, as expected, have typical heterochromatin features, *i*.*e*., high levels of H3K9me2 and DNA methylation in all sequence contexts as well as low chromatin accessibility (Fig. 3a, Extended Data Fig. 6, Supplementary Table 2). Yet, reanalysis of micrococcal nuclease (MNase) profiles (Supplementary Table 2) showed that well-positioned nucleosomes are substantially absent from ITRs except at embedded TEs (Extended Data Fig. 6). Hence, in contrast to canonical *CEN178* satellites, ITR domains do not attract nucleosomes at regularly spaced intervals, and their chromatin state differs from those of centromeres, telomeres, and TEs.

**Fig. 3.**
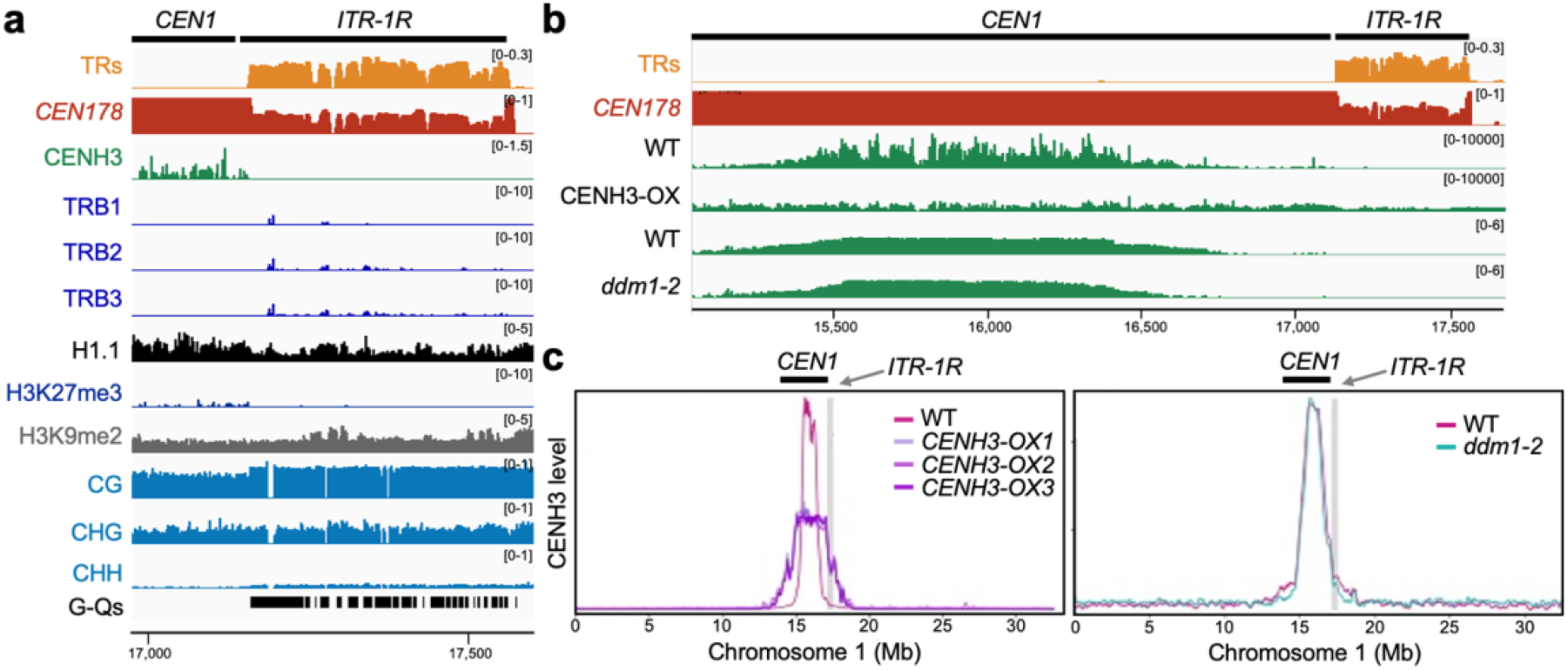
ITRs display no centromere or telomere chromatin hallmarks. **a**, Epigenome landscape of centromeric, telomeric and TE chromatin hallmarks. Datasets are detailed in Supplementary Table 2.**b**-**c**, CENH3 levels in three *CENH3* overexpressing lines (CENH3-OX) and in *ddm1-2* mutant plants are compared to their respective wild-type seed lots. The *ddm1-2* and its wild-type control ChIP signals are given as Log2(IP/Input) calculated from read counts. ChIP-seq signals of the *CENH3* overexpressing lines and cognate wild-type samples are given as RPKM.

We tested whether CENH3 ectopic accumulation could force its incorporation into ITR chromatin. Reanalysis of three independent *CENH3* overexpression lines confirmed that CENH3 spreads over the centromeres and pericentromeres^24^, and further showed this effect across the TR-rich 3’ domain of *CEN4* but significantly less at ITRs (Fig. 3b-c). To test whether CENH3 incorporation at ITRs is restricted by their heterochromatic status, we profiled endogenous CENH3 by ChIP-seq in a *ddm1* mutant line impaired in cytosine methylation^25^ and heterochromatin formation^26^. CENH3 occupancy at both ITRs and at the TR-rich *CEN4* region was similar in wild-type and *ddm1-2* plants (Fig. 3b-c), indicating that heterochromatin formation is not the main trigger of CENH3 exclusion from TR-rich *CEN178* satellites. CENH3 depletion at ITRs is consistent with their topological and functional insulation from the neighboring centromeres.

### Telomere-independent ITR size variations

To gain insight into ITR diversification mechanisms in natural contexts, we profiled the T2T genome assemblies of 110 *A. thaliana* accessions using *ITR-scan* and examined the structural diversification of all major ITR domains identified using an ITR dot-plot atlas <u>(Supplementary Video 1</u>; Supplementary Table 1, Supplementary Table 4, Extended Data Fig. 7). Long ITRs (>10 kb) were found in all the accessions examined. They are consistently made of *CenTel* satellites of similar sizes, structures, and TR density to those in Col-0, indicating that centromere-derived ITR emergence may predate *A. thaliana* diversification (Fig. 4a-e, Extended Data Fig. 7). Accordingly, *ITR-1* was universally found adjacent to *CEN1* (Fig. 4b, Supplementary Table 4). Yet, *ITR-1*/*ITR-4* sizes vary substantially across accessions, from about 200 kb to 2 Mb (Fig. 4a, Extended Data Fig. 7, Extended Data Fig. 8), suggesting that efficient mechanisms drive their expansion or contraction. *ITR-1* is typically longer than *ITR-4* and composed of one dominant (type A) or multiple HORs (type B), whereas *ITR-4* is most often shorter than 100 kb and made up of a single HOR (type C) (Fig. 4a-b, Extended Data Fig. 7, Supplementary Table 4). Against the general trend, multiple relict accessions have large (>100 kb) *ITR-4* domains with diverse HOR types (type B) (Fig. 4a, Extended Data Fig. 7-8, Supplementary Table 4). Among this group of accessions, *ITR-4* often displays a progressive decline in *CenTel* satellite density with increasing distance from *CEN4* (Fig. 4c), consistent with a gradual specification of *CenTel* sequence identity.

**Fig. 4.**
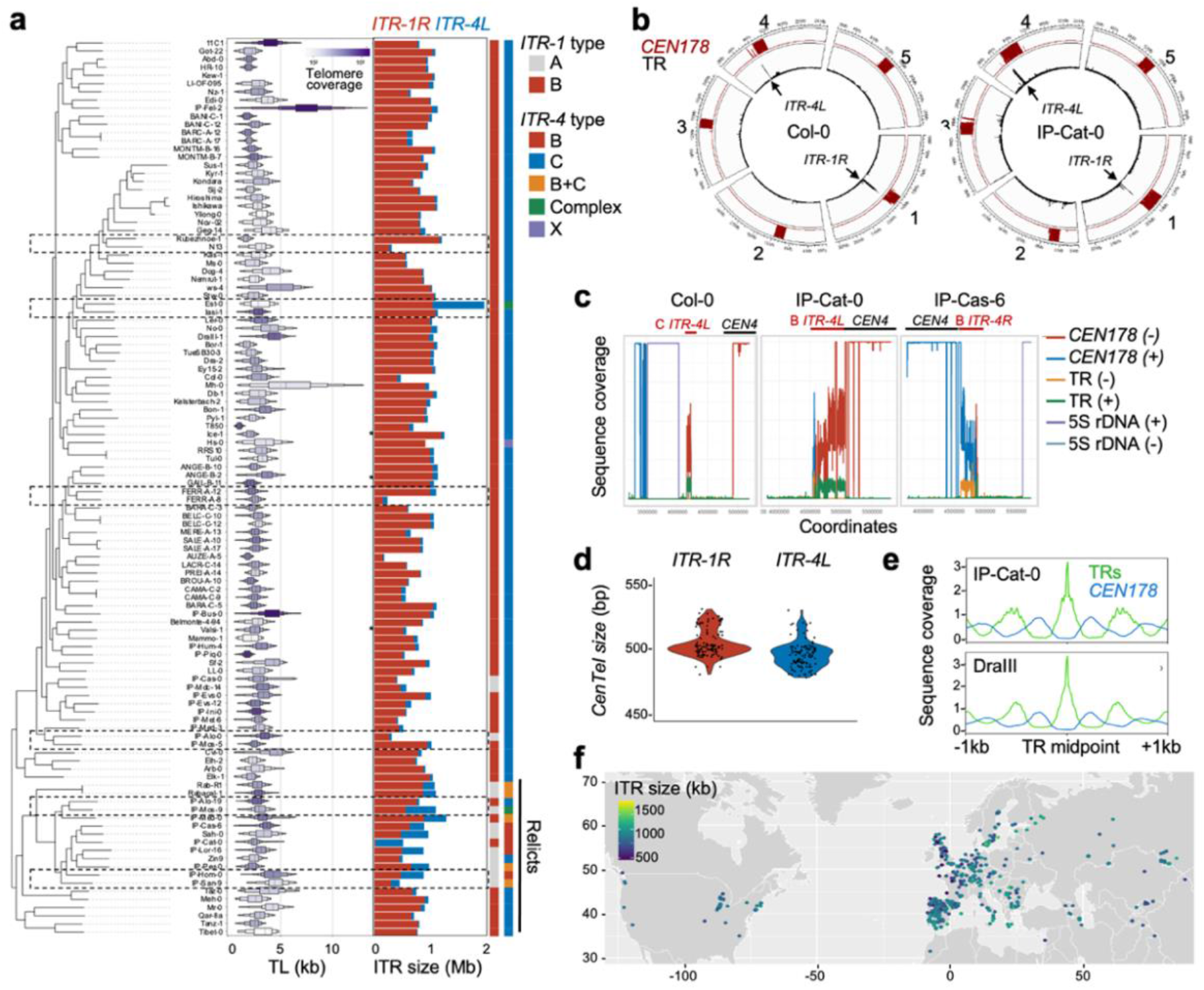
ITR content varies independently of telomere length, population structure, and environmental contexts. **a**, Phylogenetic analysis of telomere length distribution (violin plots) and ITR size (bar plots) across 110 accessions. ITR types are indicated, with boxes highlighting related accessions with substantial ITR size variations. **b**, While *ITR-4* is separated from *CEN4* by several TEs and protein-coding genes in Col-0 and other accessions, it abuts the centromere in relict accessions such as IP-Cat-0. **c**, Density of *5S* rDNA and *CEN178* satellite variants in representative *ITR-4* types. *ITR-4* is small and compact (type C) in Col-0 and large and heterogeneous (type B) in IP-Cat-0 and IP-Cas-6. In IP-Cat-0, *CEN178* density across *ITR-4* steadily declines as the distance from *CEN4* increases, with a complete inversion of *ITR-4* orientation in IP-Cas-6. **d**, Conserved average size of *CenTel* satellites in *ITR-1* (red) and *ITR-4* (blue) across the genome assemblies of 110 *A. thaliana* accessions, with one dot per accession. *CenTel* satellites are typically ∼500-bp-long with slightly more size variations in *ITR-4* than in *ITR-1* domains. **e**, Sequence coverage of TRs and *CEN178* satellites within *ITR-1* and *ITR-4* relative to each TR mid-point in two representative accessions. f, Geographical distribution of ITR size in 788 accessions estimated from NGS data of the 1001 Genomes Consortium^29^, showing no geographical representation biases.

To test whether ITR dynamics are linked to telomere diversification^27^, for example, by sharing connected regulatory forces, we used *TeloReader* to estimate telomere length across the 110 accessions (Fig. 4a, Extended Data Fig. 8, Supplementary Table 4). After benchmarking the approach using a Southern-blot-based survey of TLs in 21 accessions^28^ (Extended Data Fig. 8), we detected substantial variations in TL mean values across the 110 accessions, from 1.1 ± 0.31 kb to 7.3 ± 2.5 kb (Fig. 4a, Extended Data Fig. 8). As exemplified by IP-Piq-0 and IP-Hum-4 in Fig. 4a, a phylogenetic analysis of all accessions’ genome sequences using a set of 617,537 SNPs (excluding ITRs) showed a lack of correlation between accessions’ relatedness and mean telomere length (r^2^=-0.039; Extended Data Fig. 8). Likewise, no correlation was detected between ITR content and telomere length, consistent with both evolving rapidly and independently (Extended Data Fig. 8).

Remarkably, inspection of closely related accessions revealed sudden changes in ITR content (*e*.*g*., Est-0 *vs* Iasi-1) (Fig. 4a, Extended Data Fig. 8). We thus explored potential geographic or environmental effects on ITR content. We exploited high-quality short-read data for 788 accessions^29^ and estimated *CenTel* satellite content per genome using a k-mer approach that identifies the combined presence of telomeric and centromeric sequences within >225-bp read pairs (Extended Data Fig. 8). We benchmarked the approach by comparing 35 accessions with both Illumina and T2T genome sequences (Extended Data Fig. 8). A pangenome analysis confirmed the universal presence of substantial, yet highly variable, ITR content across accessions (ranging from 466 kb to nearly 1.6 Mb). Similarly, ITR content was uncoupled from geographic relatedness (Mantel test r=0.181, p-value=0.377; Fig. 4f, Extended Data Fig. 8, Supplementary Table 4) or local environmental conditions (Mantel test and Pearson correlation r<0.15; Extended Data Fig. 8). Collectively, these analyses reveal that ITR content and architecture diversify rapidly, often abruptly, and primarily independently of mean telomere length, population structure, and environmental contexts.

### TE-associated abrupt variations of ITRs

We searched for mechanisms behind sudden ITR size changes among related accessions by visually examining the dot-plot atlas of ITRs across the 110 accession genomes assemblies. This revealed recurrent duplications of domains spanning entire HOR blocks and adjacent TEs (Fig. 5a, Extended Data Fig. 7, <u>Supplementary Video 1</u>), which, as in Col-0, are enriched for *ENSPM* transposons (Fig. 5b, Extended Data Fig. 9). TE-associated ITR block duplications were ascertained by inspection of TE insertion sites across multiple accessions (Extended Data Fig. 9-10). In IP-Cat-0, *ITR-4L* is 98 kb longer than in IP-Med-0 due to a 100-kb HOR duplication bordered by duplicated *LINE* retrotransposons (Fig. 5a, Extended Data Fig. 10). Another striking exam-ple is *ITR-1* from the accession Tibet, which harbors 10 TE-HOR blocks duplicated in tandem and linked to the same *ENSPM* insertion event (Extended Data Fig. 9). Having found that closely related accessions repeatedly exhibit TE-HOR block duplications linked to sudden variations in ITR size and architecture, we examined at a broader scale whether TE integration and multiplication are linked to ITR expansion. Beyond the expected correlation between ITR size and the number of embedded TEs (r^2^>0.9) (Fig. 5c, Extended Data Fig. 9), we observed that ITRs exceed 100 kb only when they contain at least two similar TE copies, reaching a ∼1 Mb plateau when they carry 10 or more TE multicopies (Fig. 5d). Consistently, the ages of ITR-embedded TEs, as estimated by their Kimura distances, are on average significantly lower than those of TEs located within centromeres or elsewhere in the genome (Fig. 5e). We conclude that TEs integration mediates recurrent cycles of ITR block duplication events.

**Fig. 5.**
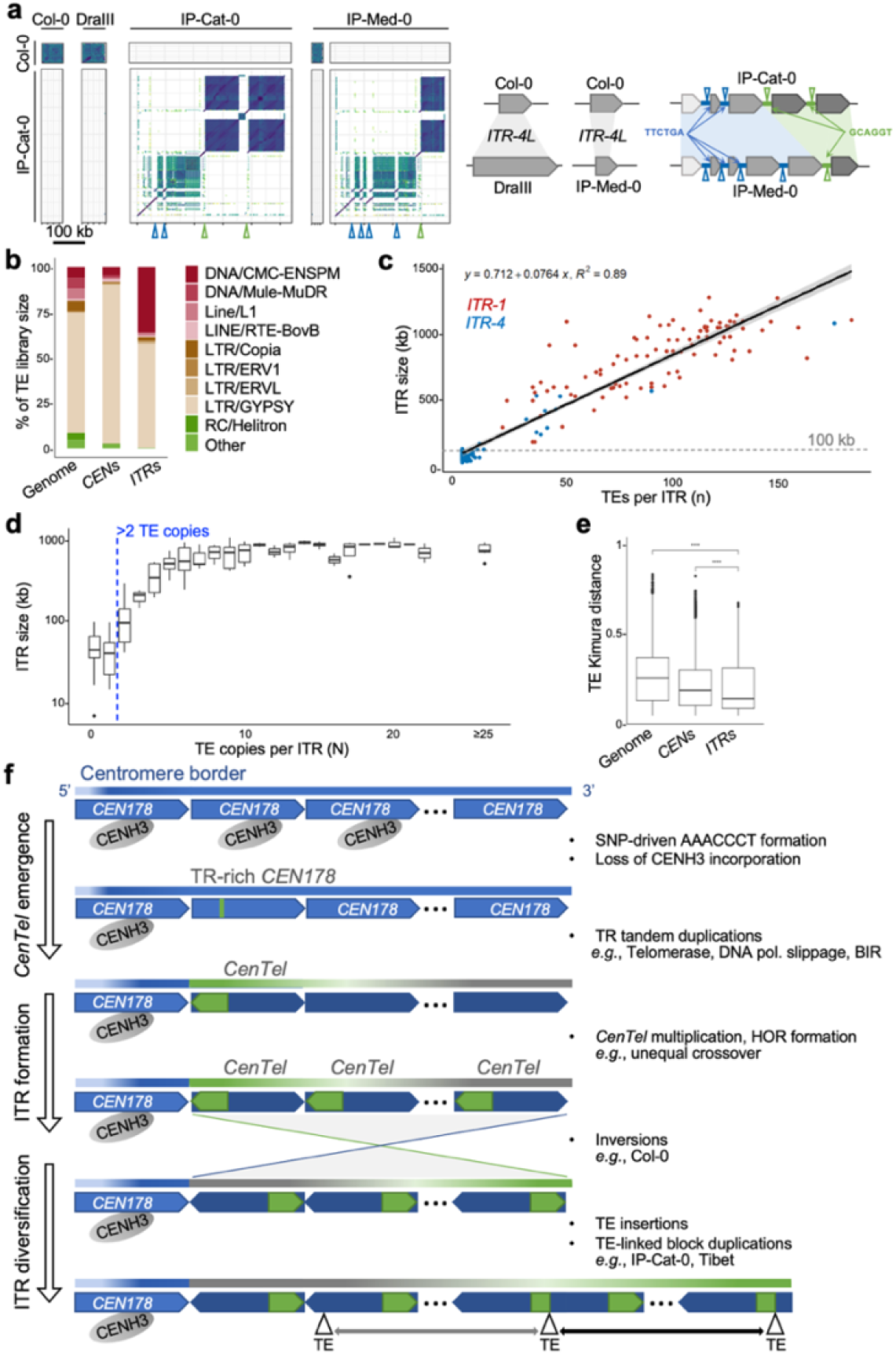
TE-associated ITR block duplications. **a**, Syntenic architectural ontions in *ITR-4L* of the non-relict Dra-III (type C) and relict IP-Cat-0 and IP-Med-0 (type B) accessions compared to Col-0 (type C) using a 50-bp sliding window. While differing in size and spacing from *CEN4, ITR-4L* domains of IP-Cat-0 and IP-Med-0 display similar HORs but differ by the duplication of large TE-associated HOR blocks. TE copies with identical integration sequences are color-coded. **b**, Distribution of all annotated TE family members in the full genomes (Genome), centromeres (*CEN*) and ITRs of 95 *A*.*thaliana* accessions, shown as the proportion of the TE family size. **c**, Comparison of ITR size and the number of embedded TEs across accessions. Dots represent individual *ITR-1* (red) or *ITR-4* (blue) domains. **d**, Comparison of maximum TE copy number and cognate ITR size. **e**, Kimura values for TEs embedded with ITRs (ITRs), centromeres (CENs) or across the genome (Genome), quantifying the sequence distance of ITR-embedded TEs from their cognate TE family consensus sequence. Willcoxon test **** p≤0.001. **f**, Evolutionary scenario summarizing our findings on ITR formation and diversification through: 1) stabilization of a *CEN178* single-nucleotide point mutation forming a telomeric motif and repelling CENH3, as found at TR-rich *CEN4* and *CEN5* domains; 2) emergence of a primordial *CenTel* satellite upon TR spreading, most likely involving other mechanisms than point mutations alone; 3) tandem duplications and homogenization of *CenTel* satellites; 4) centromere-like HOR formation^2,37^ and TE insertions promoting intra-ITR large block duplications and sudden ITR size changes.

## Discussion

Classic models view ITRs as leftover structures of telomere-associated chromosome rearrangements ^5,14,15,30^, yet these scenarios cannot readily account for the hybrid architecture of *CenTel* ITR-building satellites identified in our study. Moreover, the genomic locations of major *A. thaliana* ITRs do not coincide with the breakpoints of ancestral chromosome fusions inferred from the evolutionary history of extant Brassicaceae genomes^31^. Our data provide compelling evidence for an unappreciated centromere-oriented model in which ITRs emerge upon mutation-driven *de novo* gain of telomeric motifs, followed by satellite neo-functionalisation, local amplification, and insulation from the functional centromeres (Fig. 5f). This model of ITR formation through *in situ* sequence drift compellingly aligns with the long-noted cytological colocalization of ITRs and centromeric domains across eukaryotes^4,8^. It is also supported by our forward simulations of centromere evolution using empirical PM rates and spectra, which identified centromeres as genome-wide hotspots for spontaneous formation of telomeric motifs, and estimated that over 500 independent *CEN178* satellites can gain TRs in fewer than ten million generations. Consistently, we identified thousands of “transitional” telomeric motifs within *CEN4* and *CEN5*, altogether underscoring centromeres as major reservoirs for ITR seeding. In contrast to the propensity of centromeric satellites to form single telomeric motifs, cumulative gains of these sequences in a single satellite were found to be exceedingly rare, hinting that additional mechanisms amplify newly formed telomeric motifs to form the *Telo*-island arrays characteristic of ITRs. This presumably involves a sequence-specific route, such as telomerase-mediated ectopic healing of double-strand breaks^32,33^. As meiotic crossovers are limited at centromeres and pericentromeres^34,35^, DNA double-strand breaks may be linked to the formation of unstable G-quadruplexes or to cut-and-paste cycles of embedded TEs. Replication slippage, break-induced replication (BIR), or microhomology-mediated end joining (MMEJ) could also contribute to TR local amplification or depletion^33^. Satellite variants have long been proposed to arise upon point mutations, and to amplify and homogenize through unequal recombination and gene conversion^36^. Our PM-based simulations predicted that TR-containing *CEN178* satellites persist for hundreds of millions of generations, a temporal window that provides ample opportunities for *CenTel* emergence and fixation, *cis*-proliferation, and HOR formation via evolutionary dynamics similar to those of centromeric satellites^2,37^. Long ITRs tend to be enriched for *ENSPM* DNA transposons and *GYPSY* retrotransposons that, while obstructing HOR formation, may accelerate ITR expansion by facilitating large block duplications. Consistently, we observed that ITRs exceed ∼100 kb only when they carry multicopy TEs, which typically originate from a single integration event. Added to previous reports that TEs may transpose telomeric repeats^5^ and contribute to genome expansion^38^, TE sequences may create homology substrates for intra-ITR recombinations, thereby explaining abrupt changes in ITR size across closely related accessions, independently of geoclimatic variables or telomere length.

As established by analyses of pangenomes^35^ and of MA lines by Dong et al. (2026)^37^ as well as this study, centromeric satellites are among the fastest-evolving sequences in *A. thaliana*, yet their diversification is evolutionarily constrained by interactions with CENH3^39^. Therefore, escaping the selective pressures associated with centromere function is presumably essential for the transition of *CEN178* satellites into ITRs. Consistently, we observed that TR-containing *CEN178* variants display low CENH3 levels and that ITRs are resistant to CENH3 incorporation when forcing its spreading across neighboring pericentromeric regions or when altering ITR heterochromatic state. Given that CENH3 binding diminishes with *CEN178* consensus sequence divergence^2^ and that nucleosomes are irregularly spaced over *CenTel* satellites (this study), CENH3 association is possibly prevented by the intrinsic sequence or architectural features of the *CenTel* satellites, most likely the interruption of the satellite arrays by TR clusters or, as seen by incorporating telomeric repeats into a yeast artificial chromosome^5^, as a direct consequence of the telomeric sequence itself.

The conservation of *CenTel* satellites across all 800 accessions examined suggests that preserving a substantial ITR genome load may confer a selective advantage rather than reflecting neutral genomic turnover alone. We cannot rule out a contribution of ITRs to telomere maintenance in cells lacking telomerase activity by serving as templates for telomere elongation^27^, yet the observed independence between ITR content and telomere length variations argues against shared regulatory mechanisms with telomeres in *A. thaliana*. A homeostatic model in which ITRs act as “telomeric ballast” is appealing but difficult to reconcile with the broad ITR size changes observed across closely related accessions. Defining the mechanisms that shape ITR birth and diversification in lineages with distinct centromere and telomere architectures may help clarify the regulatory mechanisms controlling these enigmatic domains and their contributions to genome evolution.

## Acknowledgments

The authors are grateful to the TAIR12 Initiative for sharing information on the Col-CC genome assembly prior to publication, and Xiao Dong and Raúl Wijfjes (LMU, Germany) for providing the Col-CC assembly. They thank Chris Bowler (IBENS, Paris, France) for support, Simon Amiard and Aline Probst (GrED, Clermont, France) and Vincent Colot (IBENS, Paris, France) for conceptual input, PNDS and Q-lab team members for informal contributions.

## Author contributions

F.B., L.Q., and Z.X. designed the study and supervised the work. I.B. performed ChIP-seq experiments. I.B./A.E., C.G., and A.V., respectively, analyzed ITR sequences, telomere sequences, and Hi-C data. A.E. conducted the analyses of MA lines and sequence evolution simulations. I.R.H./R.B provided TE annotations across accessions’ genomes, and L.Q. provided TE annotations in the Col-CEN genome assembly. K.F., F.A.R., and D.W generated the MA lines sequencing data. F.A.R. analyzed ITR assembly quality across genome assemblies. D.D.C. provided lab management support. F.B., L.Q., Z.X., I.B., and A.E. wrote the manuscript. F.B., L.Q., I.B., C.B., and J.F. contributed to data interpretation. All authors have proofread the manuscript.

## Requests

Requests for materials and information should be addressed to Fredy Barneche and Leandro Quadrana.

