## Supplemental information for "Centromeric origin of megabase-scale *Arabidopsis* interstitial telomeric repeat domains"

### Supplementary information

### Methods

#### Estimation of telomere length from long sequencing reads

Individual sequencing reads in fastq format were collected for all accessions[2,35,40–42]. *TeloReader*[17], detecting a given telomeric motif and all its circular permutations within sliding windows, was applied to all reads. The TTTAGGG motif was used as an argument with all parameters set to default. To exclude interstitial telomere sequences, only telomere sequences longer than 500 bp and found <50 bp away from a read terminus were kept and used to plot the telomere length distributions (Extended Data Fig. 1b). For the analysis of chromosome end-specific telomere distributions shown in Fig. 1a, all reads containing terminal telomere sequences from the previous step were aligned using BLAST against the terminal 20 kb regions of all chromosome ends of the *Col-CC* genome assembly. Reads were assigned to their corresponding chromosome end according to their e-value and to the length of the alignment. This method could retrieve reads aligned to all chromosome ends except the two associated with *NOR2L* and *NOR4L*. To identify the telomeric reads associated with chromosomes *2L* and *4L*, all telomere-containing reads were aligned against one another using BLAST and a hierarchical clustering was performed based on the -Log(e-value) of all pairs, using *fcluster* with 10 clusters of approximately equal size (Extended Data Fig. 1c), and correct clustering was confirmed by visual inspection of reads within each cluster, which aligned well with one another (Extended Data Fig. 1d).

#### *ITR-Scan*

ITRs were identified in independent assemblies of the *A. thaliana* reference accession Col-0 and 112 T2T genomes of 110 *A. thaliana* accessions using the *ITR-Scan* pipeline (Supplementary Table 1, Supplementary Table 4). The 112 genomes analyzed were selected from the cited studies based on the absence of assembly gaps within or abutting the two large ITRs. As described in the accompanying *GitHub* resource (Supplementary Table 3), genomic coordinates of all frames of the canonical (TTTAGGG)n TR on both DNA strands were identified using *fastaRegexFinder*. Downstream analyses of repeat distribution were made using *bedtools* version 2.26.0[43]. All overlapping TR frames were merged using *bedtools merge* to obtain arrays of canonical TR, and first and last arrays for each chromosome, representing telomeres, were omitted from later studies. TR density was calculated using *bedtools coverage* over 5-kb windows sliding by 1-kb generated with *bedtools makewindows*. For the 110 accessions, a simplified version of this pipeline was used, calculating as described above the genomic density of only one TR frame (AAACCCT) and using a lower threshold of 0.05.

#### ITR periodicity

Each ITR was represented as a binary vector in which ones indicated the midpoints of either TRs or *CEN178* satellites. The spatial domain of each resulting binary vector was then converted to the frequency domain by employing Fast Fourier Transform (FFT), as implemented in *Numpy* v1.24.4[44]. The resulting frequencies were then used to compute the power spectrum density (PSD) by taking their absolute values and squaring them. Base-pair periodicities were obtained from the reciprocals of the computed frequency bins for FFT.

#### TE annotations and sequence analyses

Coordinates of *CEN178* and rDNA satellites were identified using *RepeatMasker* version 4.1.5[44] as described in Rabanal *et al*, (2022)[41]. Dot plots depicting ITR structure were generated using the *nucmer* option of MUMer version 3.0[45] with 50-bp windows unless stated otherwise. Metaplots generated using the *deeptools* 3.5.4 commands *computeMatrix* and *plotProfile*. To define HOR borders, monomers, and the number of units within the HOR monomer, units were defined as starting with a *Telo*-island and ending at the next *Telo*-island. Their DNA sequences were extracted using *bedtools getfasta* and compared to each other using *ClustalOmega* (Supplementary Table 3). The resulting distance matrix was plotted as a heatmap in *RStudio* version 4.3.2, with each pixel representing one unit. Similar units were visually distinguished from there, with HORs appearing as arrays of such units. G-quadruplex formation was predicted using *fastaRegexFinder*. The coordinates of non-repetitive, TE-containing ITR domains were obtained by subtracting repeat clusters from the ITR coordinates using *bedtools subtract*. By comparing ITR sequences between *Col-CC* and *Col-CEN* genomes using *nucmer*, we inferred that they correspond to identical TEs, apart from a first *ITR-1R* domain misassembled in *Col-CEN*

(Extended Data Fig. 5b). TEs were annotated in the *Col-CEN* genome using REPET (Supplementary Table 3), and across the T2T genomes of 95 accessions using the *EarlGrey* pipeline[46]. Repeats unassigned to a given TE family and TE annotations overlapping with *CenTel* arrays were excluded. TEs were defined as similar copies when clustered within the same RND family in the *EarlGrey* pipeline.

**Chamfer distances**

We computed the pairwise Levenshtein distances among all *CEN178* satellite sequences and normalized their length (dividing by the longest sequence length) to reduce size biases. We quantified the dissimilarity between genomic regions containing satellites (*CEN1-5* and *ITR-1*/*ITR-4*) using the symmetric Chamfer distance. For two genomic regions, T and S, the Chamfer distance is the average of the two following nearest-neighbor distances: $Chamfer(T,S) = \frac{1}{2}\left[\frac{1}{|T|}\sum_{t\in T} min_{s\in S}\, D(t,s) + \frac{1}{|S|}\sum_{s\in S} min_{t\in T} D(s,t)\right]$, where D(t,s) and D(s,t) are the normalized Levenshtein distances between satellites t and s. To assess statistical significance, we used bootstrap resampling (N=50): each time, satellites were resampled with replacement from the regions T and S and then used to recalculate the Chamfer distance.

**CENH3 ChIP-seq**

CENH3 ChIPs were performed on 8-day-old WT and *ddm1-2* seedlings according to[47]. Three parallel CENH3 immunoprecipitations were performed using 100 µg of chromatin with 3.5 µg of anti-CENH3 (Abcam AB72001, lot 1008198-4) coupled with Dynabeads protein-A magnetic beads (Dynabeads, Invitrogen), 20% of each sample serving as input. Co-immunoprecipitated DNA was quantified using a Qubit fluorometer using the high-sensitivity quantification kit (Thermofisher) before library preparation and sequencing at BGI (Hong-Kong).

**ChIP-seq analyses**

ChIP-seq datasets listed in Supplementary Table 2 were mapped to the *Col-CEN* genome following the pipeline available on our *GitHub* resource (Supplementary Table 3). Metaplots were made from normalized tracks using *computeMatrix* and *plotProfile* from the *deeptools* version 3.5.4 suite. Metaplots were made over ITR sequences centered around the transition between the *Telo*-island and *CEN*-island, over the centromeres centered around the midpoint of the *CEN178* satellites, and along the body of marked and unmarked genes, and pericentromeric TEs. To estimate CENH3 load per *CEN178* satellite, CENH3 and input ChIP-seq data from wild-type plants were mapped to *Col-CEN* using a stringent pipeline with *bowtie* v2.2.5 (--end-to-end, --very-sensitive, --no-mixed, --no-discordant). ChIP signal, log2(IP/Input), was computed with *bamCompare* v3.5.1 (--normalizeUsing CPM, --extendReads, --samFlagExclude 3852, --minFragmentLength 100, --maxFragmentLength 400). Per-satellite CENH3 levels were quantified as the mean ChIP signal averaged across each satellite base. Satellites were then grouped as with or without TR by intersecting the TR motif coordinates with centromeric *CEN178* satellite annotations using *bedtools* v2.30. *CEN178* within centromeres and the 1.1 Mb 3' *CEN4* region were included in these analyses. Well-positioned nucleosomes were identified as in Leduque et al. (2024)[47].

**BS-seq analyses**

Adaptor sequences were trimmed using *fastp* v0.22 and the trimmed reads were mapped to the *Col-CEN* reference genome with *Bismark* v0.24.0. Duplicate mapped reads were removed with *Picard* v2.18 (Supplementary Table 3). DNA methylation was called at each cytosine using the bismark_methylation_extractor utility with the parameters -p --CX --no_overlap. DNA methylation levels were calculated at each cytosine as the ratio #C/(#C+#T), where #C and #T represent the number of unconverted and converted read bases, respectively. Methylation levels for each CG context (CG, CHG and CHH) were stored as BED files and converted into bigwig format using *deeptools* v3.5.1.

**Hi-C analyses**

Hi-C reads of 3 biological replicates (Supplementary Table 2) were mapped on the *Col-CC* genome using the *Hi-C Pro* pipeline[49] using default parameters, but filtering out duplicate valid pairs. Raw matrices at a 1-kb bin size were loaded into R and an SCN normalization was applied to the whole matrix as in Teano et al., 2023[9]. The normalized matrices of the 3 replicates were averaged. A segment of the average matrix corresponding to a region around *ITR-1* is shown with a log10 color scale.

**Point mutation rates and spectra in G32 MA lines**

Mutation Accumulation (MA) lines (MA32-59, MA32-69, MA32-89, and MA32-119) generation 32 were grown at 23 °C under a 16 h photoperiod. High-molecular-weight DNA isolation of single 26-day-old individual plants and PacBio HiFi library construction were carried out following procedures described previously[41]. The HiFi reads were mapped to the *Col-CEN* genome using *minimap2* v2.24 with default settings. Variants were called using *DeepVariant* v1.9.0 with default parameters. Multiallelic sites were decomposed into biallelic and left-normalized against the *Col-CEN* genome using *bcftools* v1.21. Variants shared by at least two lines were removed as likely inherited from the common parent. To reduce potential false positives, we applied the following filters with *bcftools*: GT="1/1" && FORMAT/GQ>=20 && FORMAT/AD[0:0]==0 && (FORMAT/AD[0:0]+FORMAT/AD[0:1]) >= 50. Only homozygous non-reference genotypes were retained for downstream analyses. PMs were used to estimate per-generation, per-PM rates, for different genome compartments as $\mu$=num_PM / (num_generations × num_lines × region_length).

**Simulation of PM-based TR emergence**

Per-compartment PMs were used to infer base-substitution spectra. To estimate the probability of TR appearance in *CEN178* sequences, we modeled the evolution of the canonical *CEN178* sequence as a discrete-time Markov chain parameterized by the estimated PM rate across *CEN178* satellites and the base-substitution spectrum estimated from centromeric regions. Concretely, the probability that a telomeric motif arises at a genomic position j, with base $x_j$ (A, C, G or T), was defined as a function of the point-mutation rate $\mu_j$ and a 4×4 base-substitution spectrum S. This function estimates the probability that the base $x_j$ transitions to $y_j$ as: $P(y_j \vee x_j, \mu_j, S) = (1-\mu_j)1[y_j = x_j] + \mu_j S_{x_j \to y_j}$,where 1[.] is the indicator function. For a k-mer TR motif m=($y_0,\ldots,y_{k-1}$) evaluated on the genomic region ($x_j,\ldots,x_{j+k-1}$), the probability that the region transitions to m was calculated using the centromeric PM rate ($\mu_j$=4.7×$10^{-8}$) as:

$$P(matj) = \prod_{i=0}^{k-1} p(y_j \vee x_{j+i}, \mu_{j+i}, S).$$

A 4x4 base-substitution spectrum S was defined from the centromeric PMs, with each row normalized to sum one and the diagonal entries set to zero ($S_{x\text{->}x}$= 0). As the k-mer TR motif, we used all seven phase rotations of the 7-bp canonical telomeric motif and their reverse complements. Prior to computing transition probabilities, we masked genomic positions already matching any phase of the AAACCCT motif to avoid counting existing TR motifs.

To estimate TR emergence rate per *CEN178* satellite per generation, we extracted all TR-free 178-bp *CEN178* sequences from the *Col-CEN* assembly and removed duplicates (n=17,233). Each satellite was evolved for t generations using the discrete-time Markov chain parameterized by the centromeric spectrum S and position-specific PM rates ($\mu_j$). Under this model, the evolved satellite sequence at generation t is represented as per-position base probabilities (178 positions x 4 bases). Using these probabilities, we computed the probability that the evolved satellite contains exactly N occurrences of a specific TR at generation t, P(c=N at t), by applying dynamic programming over an Aho-Corasick automaton. TR occurrences were tracked from 0 to N, with N+1 occurrences aggregated into a single N+1 bin to keep the state space finite. The probability P(c=N at t) was calculated independently for each of the seven phase rotations of the 7-bp canonical TR, and these probabilities were summed to obtain the probability that the evolved satellite contains exactly N occurrences of any TR at generation t. Using eight increasing time points (t=$10^0$, $10^1$, up to $10^7$ generations), we obtained 17,233 x 8 estimates that were used to compute the per-satellite per-generation probability of containing exactly N TR occurrences. In all calculations, we used PM rates estimated per satellite position from PMs identified across centromeric 178-bp *CEN178* sequences, assuming a rate of $10^{-10}$ for positions lacking PMs. As a control, each TR-free *CEN178* sequence was randomly shuffled, while avoiding the creation of TRs, and evolved for t generations under the same model parameters.

**Estimation of ITR abundance from short sequencing reads**

To infer the abundance of ITR domains in *A. thaliana* accessions using short-read DNA sequencing data of the 1001 Genomes Consortium[29], we developed a k-mer-based method. ITRs from the *Col-CEN* assembly were screened to identify regions enriched with TRs (*Telo*-islands) and flanked by sequences lacking these motifs.

Nucleotide sequences overlapping with *Telo*-islands were recovered, and their 3-mers were extracted. These 3-mers were then used to construct a forward Telo profile, which consisted of a vector containing the normalized frequencies of the possible 4^3-mers. Additionally, the reverse-complemented 3-mers were used to define a reverse Telo profile. These profiles were subsequently employed to identify ITR-derived reads. Given paired-end DNA sequencing data, each read pair was processed iteratively. Each read in a pair was represented as a vector of normalized frequencies of its 3-mers. This vector representation was then used to determine if the read was ITR-derived by measuring how well it matches the Telo profiles. The level of matching was assessed by calculating the dot product between the read vector representation and each TR profile built from the 3-mers as described above. Dot products with values higher than 0.08 were considered as matches, and read pairs were considered ITR-derived if exactly one read matched any of the Telo profiles. The threshold value was determined by maximizing the Pearson correlation coefficient between the ITR lengths observed in 35 assembled *A. thaliana* accessions and those estimated by the k-mer-based method. The number of read pairs matching the Telo profiles was used as a proxy for ITR length. To avoid artifacts arising from heterogeneous sequencing depths across *A. thaliana* accessions, both unmatched and matched reads were aligned to protein-coding gene exons in the *TAIR10* assembly. The average read depth of this mapping was then used to normalize the number of read pairs matching the Telo profile. To express estimated ITR lengths in base pairs, we fitted a linear model to regress the ITR lengths estimated from short sequencing data against their corresponding lengths observed in the assembled genomes. To improve estimation accuracy, we excluded *A. thaliana* accessions with low-quality DNA sequencing data: single-end sequencing or mean fragment sizes below 225 bp.

#### *CEN178* classification

To classify the *CEN178* satellites within ITRs, we first extracted their nucleotide sequences and then assessed their similarity using pairwise mapping with *minimap2* v2.24. The resulting similarity values were subjected to hierarchical clustering using Ward's linkage method, implemented in *SciPy* (Supplementary Table 3). The number of clusters was then automatically determined by using the *maxclust* thresholding provided by the *SciPy fcluster* function. To determine structural organization patterns among clusterized satellites, *ITR-1R* and *ITR-4L* were both represented solely by their *CEN178* sequences belonging to a particular cluster, and their pairwise sequence similarity values, obtained by *minimap2* mappings, were visualized as a heatmap.

#### Phylogeny of accession genomes

To estimate the divergence time among *A. thaliana* accessions, we inferred a phylogenetic tree using fourfold degenerate SNPs identified from their genome assemblies, including the assembly of *Arabidopsis lyrata* as an outgroup. To this end, error-free, paired-end reads were generated from each genome assembly by using *art_illumina* v2.5.8 with the following parameters: -ss HS25 -l 150 -f 20 -m 350 -s 50. Reads were mapped to the *TAIR10* assembly, and per-accession BAM files were built. SNPs were called from the BAM files by using *bcftools mpileup* v1.13. Fourfold SNPs were identified with *SnpEff* v5.2e and retrieved using *bcftools view* -i 'INFO/ANN~ "synonymous-variant"' to define a multiple sequence alignment with *vcf2phylip.py*. The resulting alignment was used for phylogenetic inference with *iptree2* v2.2.0 using a GTR model and 1,000 bootstraps.

#### ITR size correlation tests

We tested the correlation between the ITR size estimated from the NGS data and each of the 196 quantitative environmental factors described[50], while correcting for the phylogenetic distance determined in this study, using the Mantel test from the *ecodist R* package. The analysis included all 531 accessions for which data were available. Correlations between ITR size estimates and phylogenetic distances (excluding the environmental effect) were additionally tested using the Mantel test, and correlations between ITR size estimates and each of the environmental parameters, without the correction for the phylogenetic distance, were calculated using a Pearson test.

#### Data availability:

All data and materials are available in the manuscript or the supplementary materials. Other datasets and bioinformatic sources are detailed in Supplementary Tables 1-4. ChIP sequencing data generated in this study have been deposited at Gene Expression Omnibus under accession GSE317061. PacBio HiFi reads for the four MA lines (generation 32) have been deposited at the European Nucleotide Archive (ENA) under Accession PRJEB77512.

**Code availability**
Code availability is detailed in Supplementary Table 3.

**Complementary references**

**Extended data figures and legends**

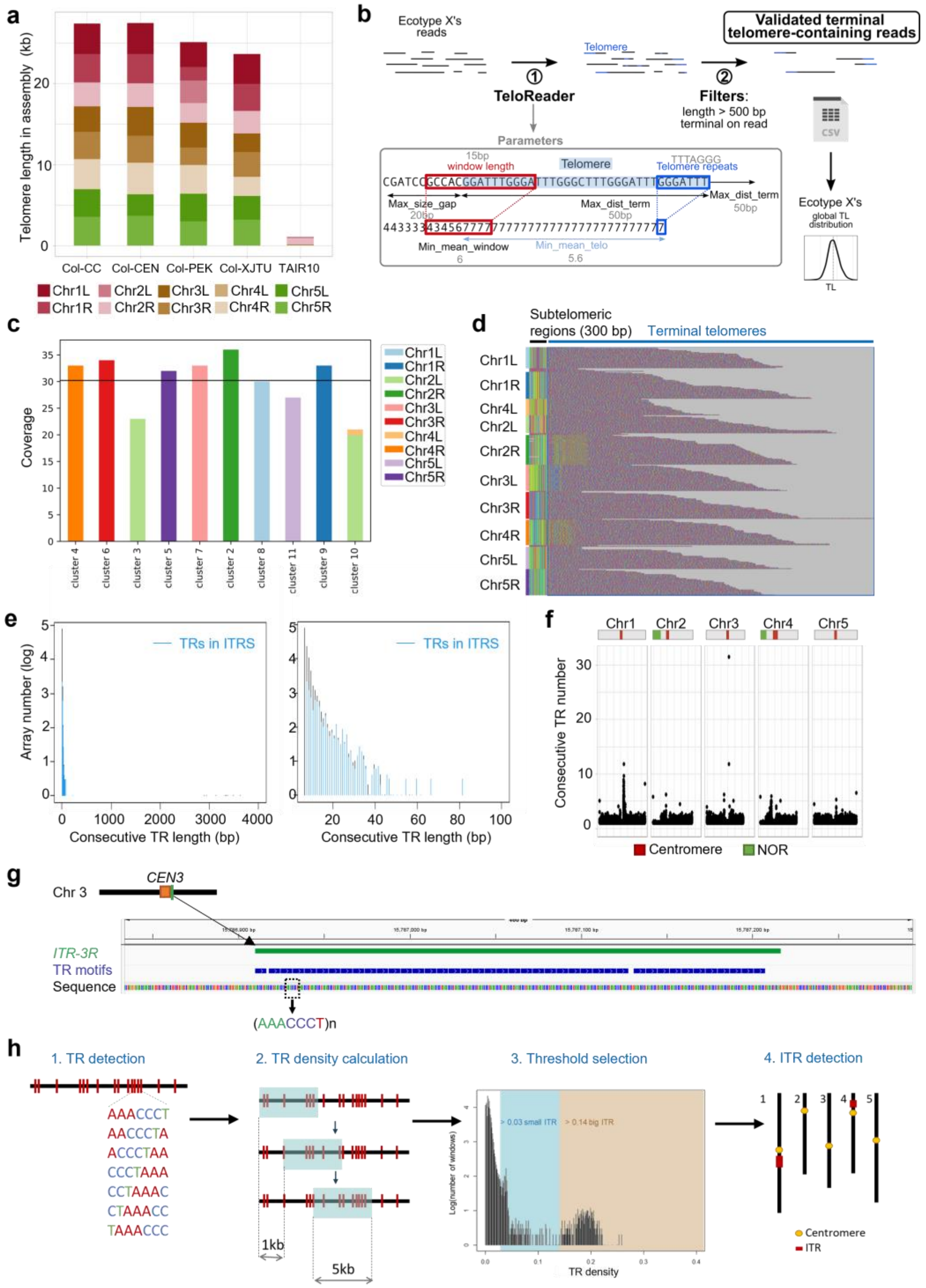


**Extended Data Fig. 1. | Telomere and ITR detection in Col-0 genome assemblies using *TeloReader* and *ITRScan*.**

**a**, Telomere length (TL) in Col-0 genome assemblies was calculated as the length of terminal uninterrupted AAACCCT/AGGGTTT arrays. A similar mean length of 3.4-kb was found in the *Col-CC* and *Col-CEN* assemblies, but shorter ones in the *Col-PEK* and *Col-XJTU* assemblies. No comprehensive data could be retrieved from *TAIR10*. **b**, Bioinformatic pipeline used to compute TL distribution from individual long-read sequences. 1) *TeloReader* was used to identify all telomere sequences in ONT or PacBio reads. The inset shows how *TeloReader* calculates a score based on the similarity to any permutation of the consensus telomere motif over a sliding window and using thresholds for the parameters (see Methods). 2) To filter out interstitial telomere sequences, only terminal telomere sequences longer than 500 bp were kept. **c**, Number of terminal telomere-containing reads in *Col-CEN* sequencing data at each chromosome extremity after alignment to the *Col-CC* genome assembly, followed by clustering analysis. The alignment step initially misassigned many *Chr4L* reads to *Chr2L*, because both extremities contain NORs. The clustering step allowed their correct reassignment to *Chr4L* (labelled as cluster 10). A black horizontal line denotes the average coverage of all clusters. **d**, Visualization in *Jalview* (with default color scheme) of all reads from the 10 clusters in **a** aligned together for each extremity, with 300 bp of nearly perfectly aligned subtelomere sequences shown. **e**, Size distribution of all TR arrays in the *Col-CC* genome assembly (black) and ITRs (blue), showing that, even in long ITRs, most TRs are not present as long continuous arrays. TRs were identified by *fastaRegexFinder* as all frames and both orientations of the consensus (AAACCCT)n motif and merged into non-overlapping arrays by *bedtools merge*. **f**, Size of continuously concatenated TR arrays divided by the TR motif size (7 bp) across the *Col-CC* genome. Regions with a large number of consecutive repeats include subtelomeric regions, small ITRs primarily present at pericentromeric regions of all chromosomes, and *ITR-1R* and *ITR-4L*. For simplicity, terminal telomeres are excluded from the plot. **g**, *ITR-3R* sequence organization. *ITR-3R* is an unusually long stretch of continuously concatenated telomeric repeats (307 bp) made out of two clusters of 30 and 10 perfect TR motifs, respectively, separated by a single imperfect TR motif. **h**, Methodological determination of TR density along the *Col-CC* genome. To identify TR-rich regions beyond continuously concatenated TRs, the *ITR-Scan* pipeline based on TR density was implemented. It divides the genome into 5-kb sliding windows with a 1-kb step, and for each window, provides the density of TR motifs as a number of bases belonging to telomeric motifs divided by the window size. Here, we plotted the distribution of TR density across all windows. Many windows expectedly display a TR density near 0, but there is a notable enrichment of windows with TR densities above 0.03 and 0.14, indicating large populations of TR-rich windows. Regions of the genome with windows above 0.14 were defined as large ITRs and regions with windows above 0.03 as small ITRs. In total, this identified 38 small ITRs in the *Col-CC* genome, predominantly near the centromeres, with varying size and structure.

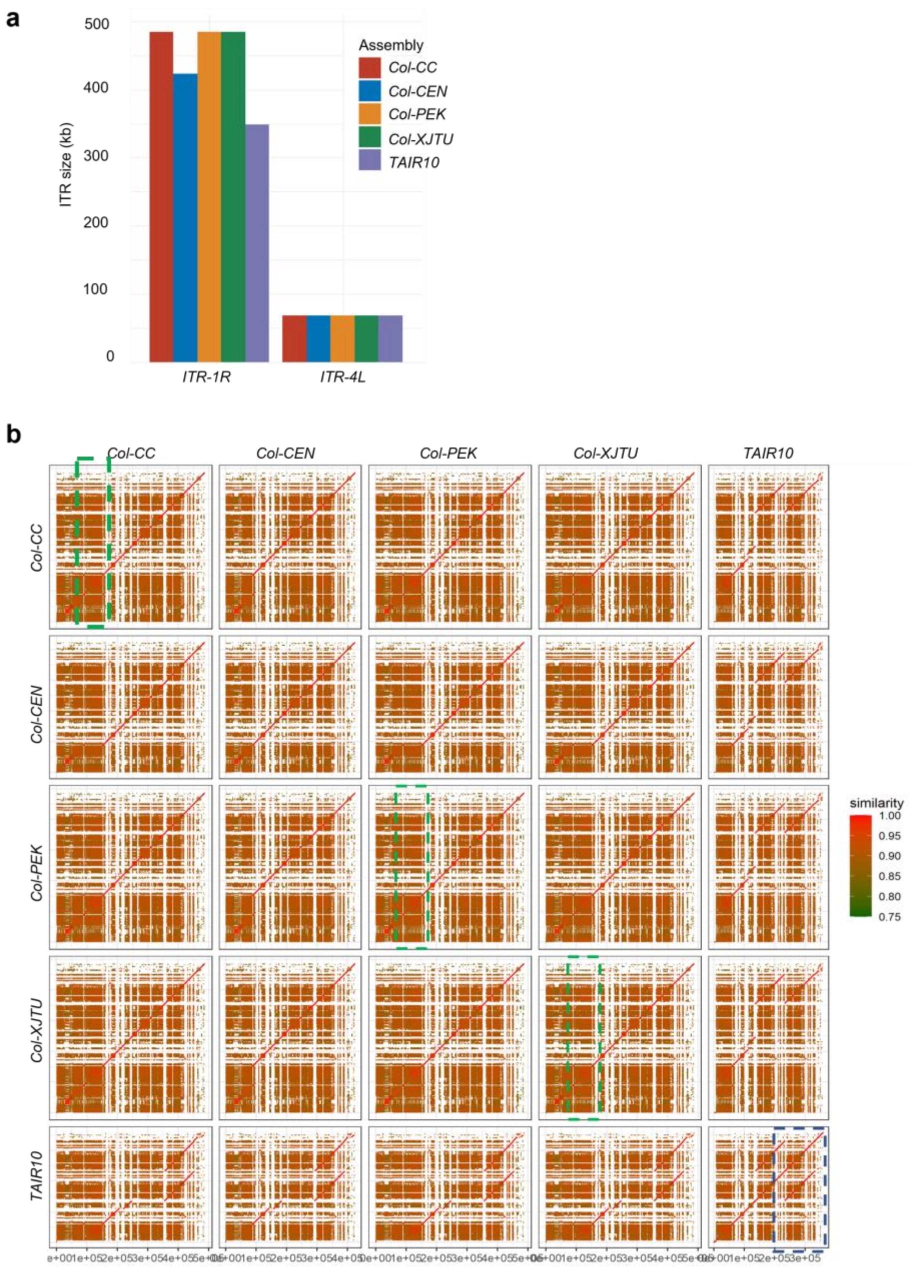


**Extended Data Fig. 2. | *ITR-1R* and *ITR-4L* are conserved across independent genome sequencing data.**
**a**, *ITR-1R* and *ITR-4L* sizes in independent Col-0 genome assemblies. In *TAIR10*, *ITR-1R* is significantly smaller than in all long-read-based genome assemblies. The real size difference is misleading since a 100-kb duplication in *TAIR10* artificially extends its size, so *Col-CC ITR-1R* and *TAIR10 ITR-1R* only share a 240-kb-long sequence. There is a 60-kb difference between *ITR-1R* of *Col-CEN* and those of the other long-read-derived genomes due to the duplication of one repeat block and a cluster of TEs not present in *Col-CEN*. *ITR-4L* length is similar in all assemblies. **b**, Sequence identity dot plot of *ITR-1R*. A duplicated block absent in *Col-CEN* is boxed in green. A block artifactually duplicated in *TAIR10* is boxed in blue. Pairwise sequence comparisons of ITRs used a 50-bp sliding window, after which the resulting similarity matrix is visualized as a heatmap, effectively creating a dot plot. Conversely, there are substantial discrepancies in *ITR-1R* length and sequence between *TAIR10* and long-read-derived assemblies, highlighting the importance of assembly curation for accurate assembly of long ITRs. Given its long size, its reproduced assembly in *Col-PEK* and *Col-XJTU*, and future annotation in TAIR12 (ms *in preparation*), *Col-CC* was used as a reference in this study, while *Col-CEN*, for which gene and TE annotations were available[2] ( Edera, Larue et al., *submitted*) was used for complementary analyses.

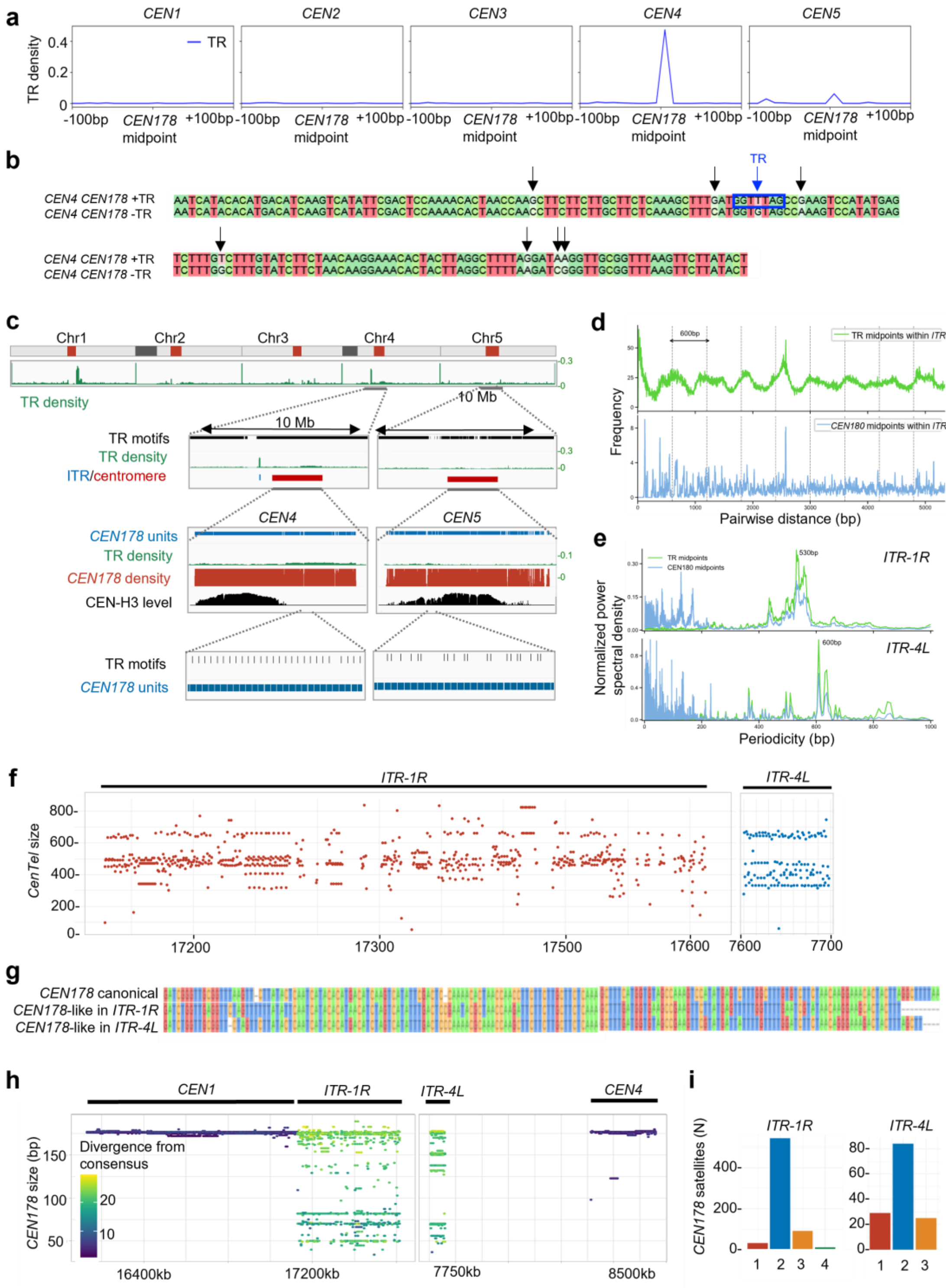


**Extended Data Fig. 3. | TR-rich *CEN178* centromeric and ITR satellites.**
**a**, TR density at centromeres centered around the mid-point of *CEN178* satellites. The coordinates of *CEN178* satellites were identified in the *Col-CC* assembly using *RepeatMasker*. As in other studies[2] centromeres were manually defined by the presence of consecutive *CEN178* satellites, rather than by CENH3 occupancy. The most common mutation is located near the *CEN178* mid-point. **b**, Sequence alignment of *CEN178* satellites from the 5' and 3' half of *CEN4*, made by *ClustalOmega*, shows multiple SNPs (arrows), including the point mutation that led to a TR (boxed in blue). **c**, *Col-CC* genome profile of *CEN178*, TR and CENH3 occupancy. An unusually high TR density characterises large domains of *CEN4* and *CEN5* after the formation of one or more TRs in *CEN178* satellites by point mutations. These mutations coincide with a sharp decrease in CENH3 occupancy.

**d**, Frequency of pairwise distances between TR midpoints (top) and *CEN178* satellite midpoints (bottom) in *ITR-4L*. **e**, Power spectral density (PSD) of the periodicity (bp) observed between TR midpoints of and *CEN178* satellite midpoints in *ITR-1R* (top) and *ITR-4L* (bottom). For each ITR, the PSD values for TRs and *CEN178* satellites were normalized so that their sums equal 1. The dominant periodicity (highest PSD value) for each ITR is indicated. **f**, Conserved size of the *CenTel* units across the ITR-IR (red) and *ITR-4L* (blue). Each unit was defined as starting with the cluster of canonical and degenerate TR-s (*Telo*-island) and ending with the last *CEN178* satellite of the *CEN*-island. , **g**, Alignment of the consensus *CEN178* sequence[2] with the consensus *ITR-1R* and *ITR-4L CEN178*-related sequences created using *ClustalOmega*. *CEN178*-related sequences were identified using *RepeatMasker* with the consensus *CEN178* sequence defined in[2] as the custom library. Their consensus was obtained using a Hidden-Markov model from their *muscle* sequence alignment. **h**, Sequence divergence of *CEN178*-related sequences in large ITRs and their proximal centromeres from the consensus *CEN178* satellite sequence, as calculated using *RepeatMasker*, showing a clear distinction between the two satellite types. **i**, Frequency of the *CEN178*-related satellites per *CenTel* unit shown as the number of *CenTel* units with the specific (1-4) number of *CEN178*-related satellites. The majority of the *CenTel* units in both *ITR-1R* and the *ITR-4L* display two *CEN178*-related satellites.

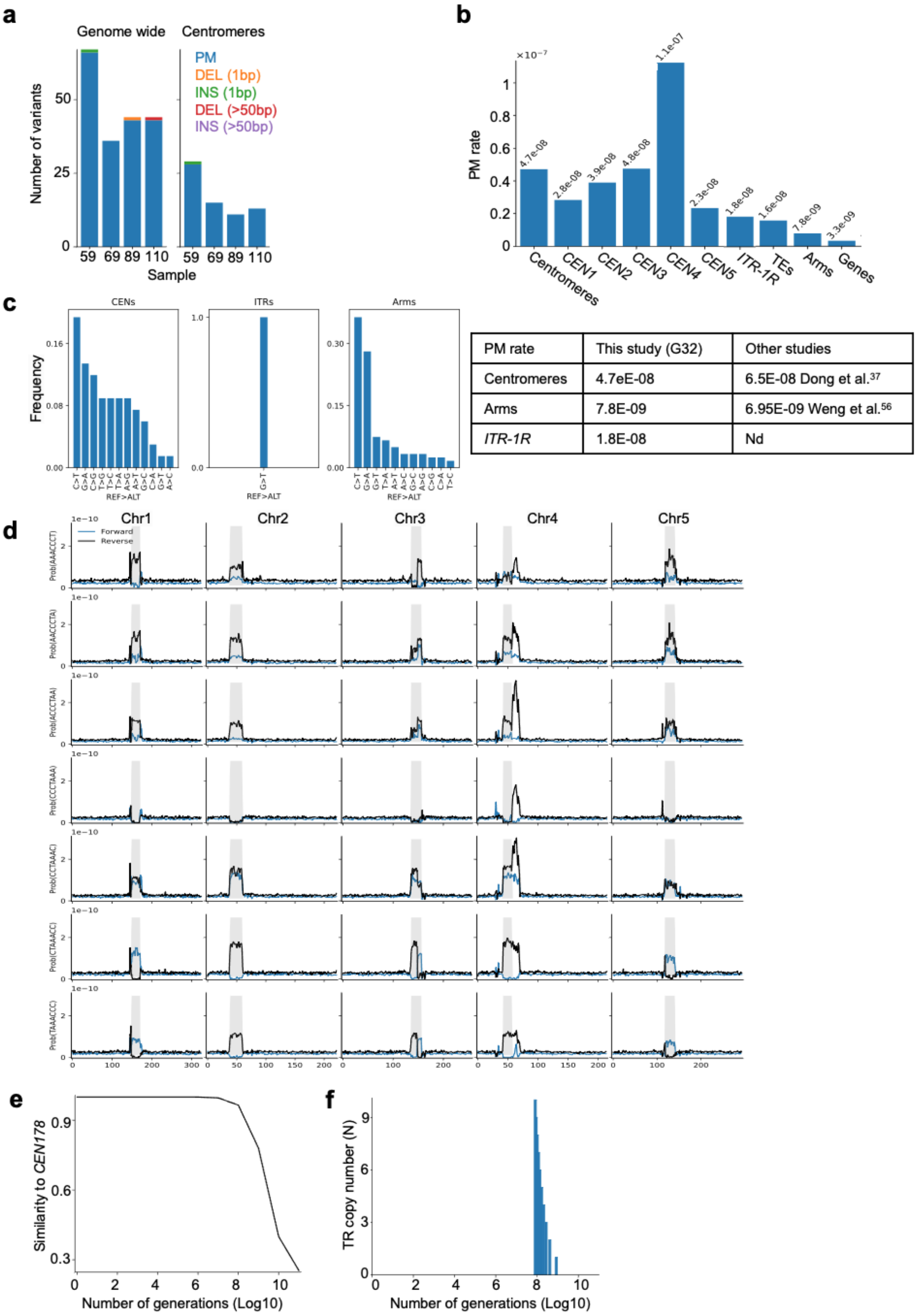


| PM rate | This study (G32) | Other studies |
|---|---|---|
| Centromeres | 4.7E-08 | 6.5E-08 Dong et al.[37] |
| Arms | 7.8E-09 | 6.95E-09 Weng et al.[56] |
| *ITR-1R* | 1.8E-08 | Nd |

**Extended Data Fig. 4. | Estimations of point mutation rates to build the *CEN178* evolution model.**
**a**, Sequence polymorphisms identified across four generation-32 Col-0 MA lines[23]. Bar plots show the number of variants identified in each MA line for each type. The left panel shows variants identified genome-wide, while the right panel shows variants identified within centromeric regions. PM, point mutation; Del, deletion; INS, insertion. **b**, Per-generation PM rates across different genomic regions identified in this study compared with those in[37,57]. Nd, not determined. **c**, Base-substitution spectrum for centromeric regions (*CENs*), *ITR-1R* (*ITRs*) and chromosome arms (Arms). **d**, Genome-wide probability of TR emergence estimated for each chromosome. Plots show the aggregated probability that any of the 7 phase shifts of the canonical TR motif (AAACCCT) appears at a given genomic position, shown separately for the forward (blue) and reverse (black) orientations. Centromere positions are indicated in gray. **e**, Modeling of the canonical *CEN178* sequence evolution using a discrete-time Markov chain parameterized by per-generation PM rate across *CEN178* satellites and the centromeric base-substitution spectrum. The canonical *CEN178* sequence is predicted to diverge substantially after ~8M generations, indicating that *CEN178* sequence identity is well conserved over evolutionary timescales shorter than speciation. **f**, The expected lifetime of a single TR within the canonical *CEN178* sequence is about 8M generations.

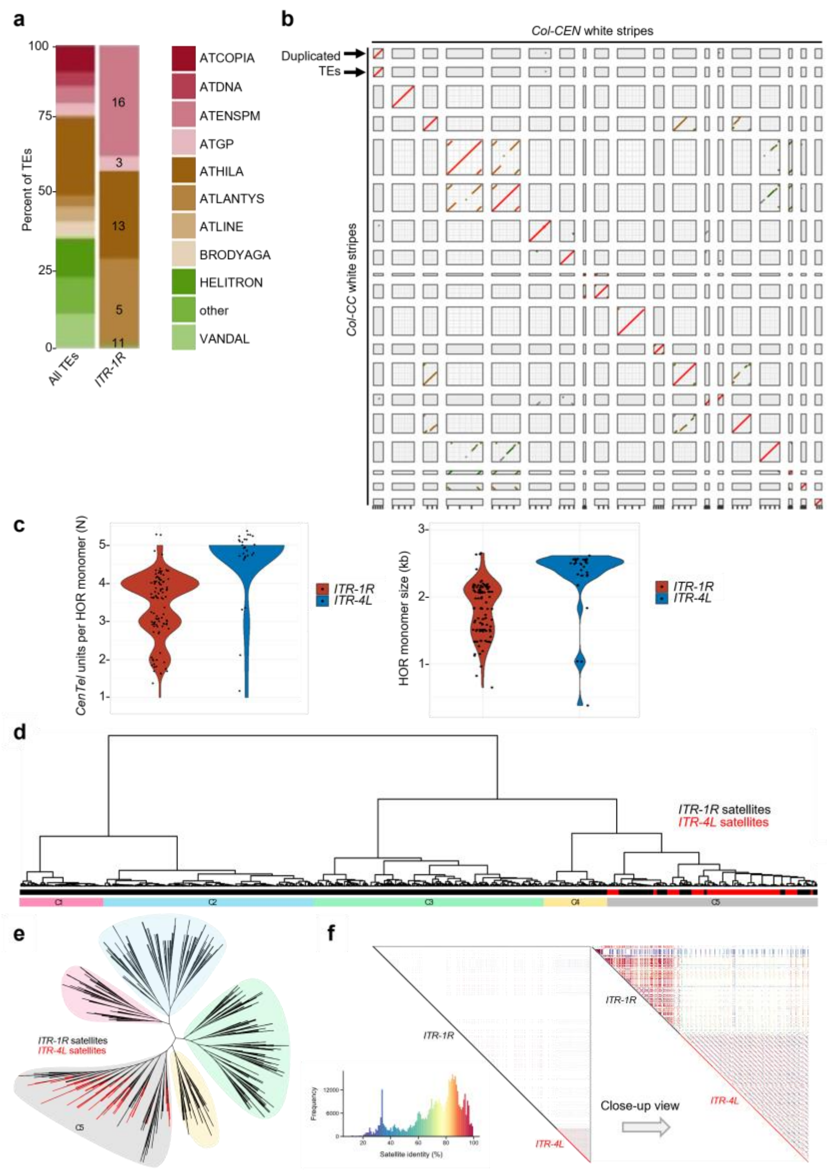


**Extended Data Fig. 5. | ITR satellite variants, HOR organization, and embedded TEs in Col-0.**
**a**, Over-representation of *ATENSPM*, *ATHILA* and *ATLANTYS* TE family members in *ITR-1R* compared to their genome-wide occurrence plotted as the percent of total TE family size. The number of TEs of each family in *ITR-1R* is given. **b**, Conservation of TE positions in *ITR-1R* between the *Col-CEN* and *Col-CC* genome sequences. TEs are first identified as poorly repeated sequences visualized as white stripes in dot-plot sequence comparisons. White stripes were extracted by subtracting clusters of repeats from the ITR coordinates using *bedtools subtract*, assembled and used for dot-plot comparisons of *Col-CC* and *Col-CEN* genome assemblies using *nucmer*. Since all but one duplicated white stripe is shared between *Col-CEN* and *Col-CC*, we used the *Col-CEN* annotation to infer TE positions in the *Col-CC* genome. **c**, Number of *CenTel* satellites per HOR monomer (left) and size of HOR monomers (right). *ITR-4L* is composed of a single HOR made out of 5 *CenTel* cluster units. Only a few HOR monomers are truncated, comprising a small number of *CenTels*. *ITR-1R* is predominantly composed of a 4-*CenTel* HOR interrupted by several 2- or 3-*CenTel* HORs. Since *CenTel* satellites are ~500 bp long, the 5-unit *ITR-4L* HOR monomers are around 2.5-kb long, while 3 or 4-unit *ITR-1* HOR monomers are 1.5-to-2-kb-long. **d**, Sequence similarity-based hierarchical clustering of *ITR-1R* and *ITR-4L* *CEN178* variants. The horizontal colored bars indicate satellite origin (*ITR-1R*/black, *ITR-4L*/red) and their clustering into five groups (C1-C5). **e**, Unrooted cladogram highlighting the close relationship between *CEN178* satellites from *ITR-1R* and *ITR-4L* within the C5 group. **f**, Bottom left panel, sequence similarities between *CEN178* satellites within *ITR-1R* and *ITR-4L*, with sequence similarity values color-coded. The white color indicates no detectable similarity. Right panel, sequence similarities between *CEN178*-related satellites across *ITR-4L*.

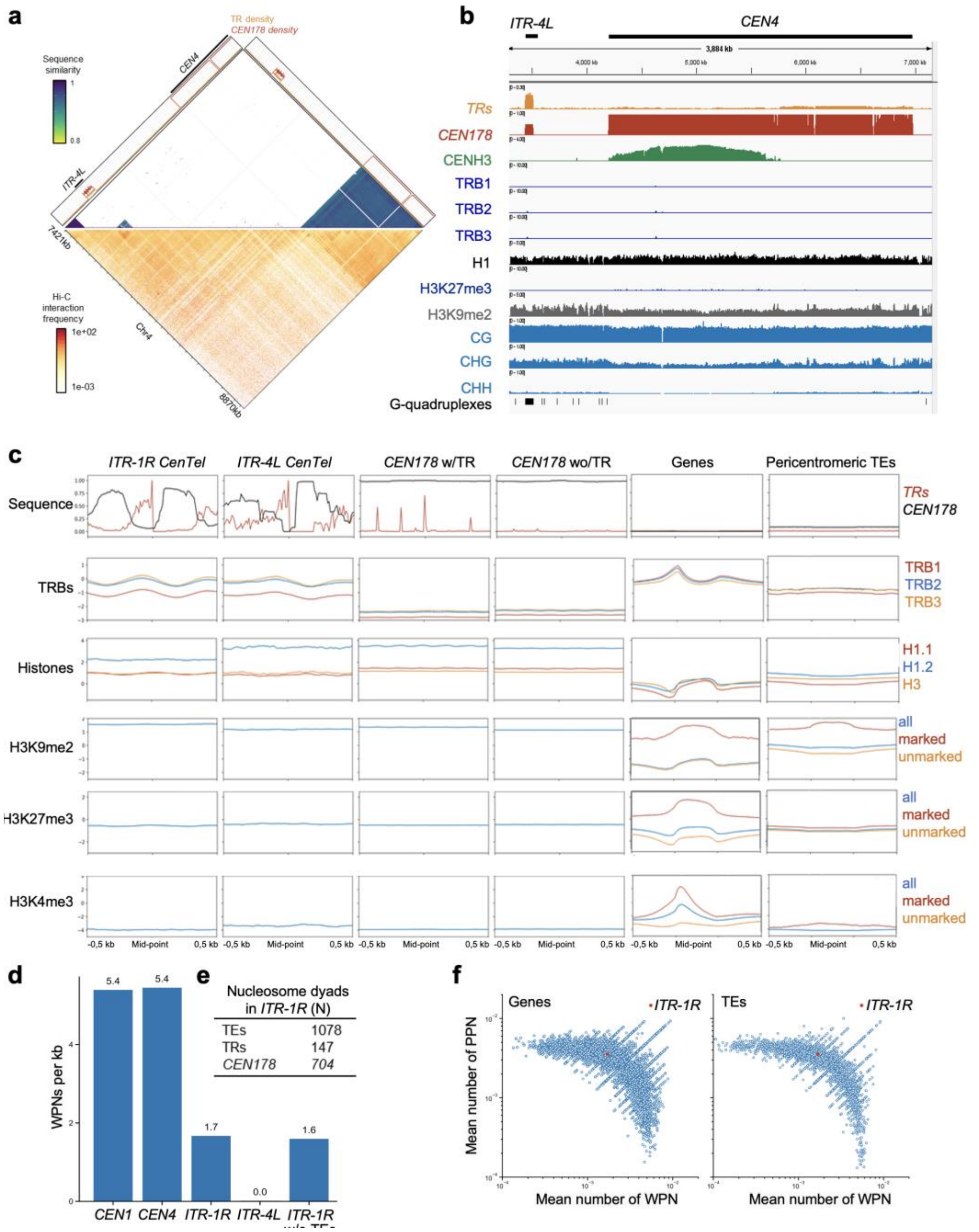


**Extended Data Fig. 6. | *ITR-4L* sequence architecture, 3D organization and chromatin properties in the accession Col-0.**

**a**, Dot plots showing sequence identity (up) and the 1-kb resolution Hi-C profile (bottom) of the *ITR-4L* region, including the left domain of *CEN4*. In the upper panel, the densities of TRs and *CEN178*-related repeats are shown in orange and red, respectively. **b**, IGV browser view of centromeric, telomeric and TE chromatin hallmarks in *ITR-4L* and *CEN4*. The corresponding datasets are detailed in Supplementary Table 2. G-quadruplex distribution was predicted using *fastaRegexFinder*. **c**, Mean enrichment of the indicated chromatin factors and histone modifications. The y-axes are scaled to compare *ITR-1R* and *ITR-4L*, distinguishing the bulk of *CEN178* satellites displaying (w/TR) or not (wo/TR), as well as to characteristic reference loci (protein-coding genes, and pericentromeric TEs). **d**, Nucleosome positioning[48] at *ITR-1R* and *ITR-4L*. Bars depict the number of well-positioned nucleosomes (WPNs) per kb within *CEN1*, *CEN4*, *ITR-1R* and *ITR-4L*, with or without considering their embedded TEs (without TEs). **e**, Number of nucleosome dyads overlapping TEs, TRs and *CEN178* satellites in *ITR-1R*. **f**, Comparison of well-positioned nucleosome (WPN) and poorly-positioned nucleosome (PPN) average number between genes, TEs, and *ITR-1R*.

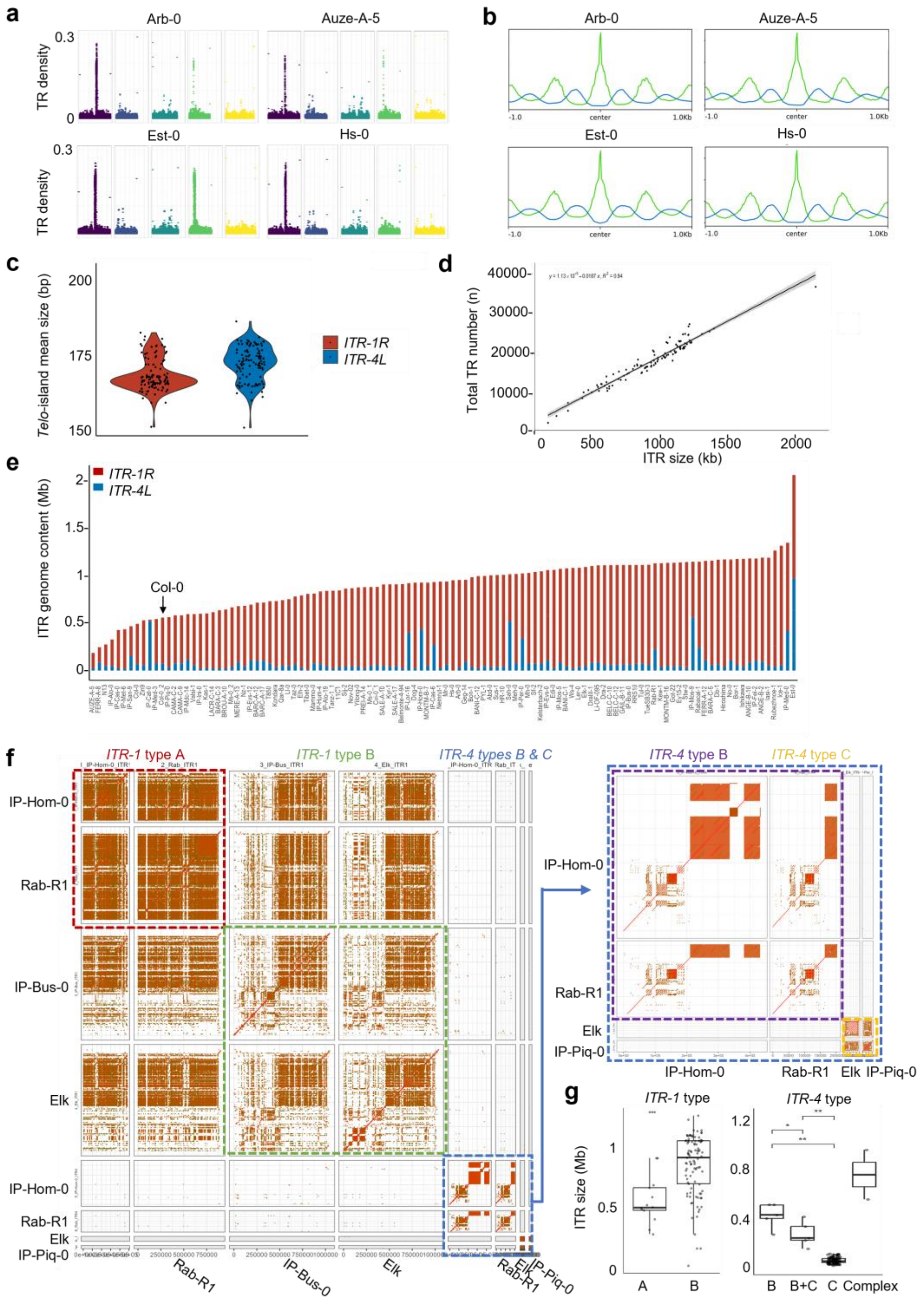


**Extended Data Fig. 7. | Conserved satellite but diverse higher ITR organizations across the T2T genomes of 110 accessions.**

**a**, TR density across the chromosomes of representative accessions. TR density was calculated as previously described for Col-0, analyzing all frames and strands of the canonical AAACCCT motif detected using *fastaRegexFinder*. TR density across genomes was calculated using *bedtools coverage* with 5-kb windows, sliding by 1 kb. Each point represents TR density in a single window. Terminal windows, representing telomeres, are excluded to better visualize the lower TR density in ITRs. TR-density profiles are conserved across the accessions, with moderate TR-increase present in the majority of the pericentromeres (small ITRs) and major TR density peaks in the pericentromeres of chromosomes 1 and 4 (*ITR-1* and *ITR-4*). Differences in the number of dots per peak were used as a proxy for ITR size differences. While *ITR-1* was found in all examined accessions, *ITR-4* was not detected in the Hs-0 accession. **b**, Meta-profiles of TR and *CEN178*-related repeats within large ITRs across all the accessions analyzed, illustrating the remarkable conservation of *CenTel* repeats with a ~500-bp periodicity. Metaplots were centered around TR using *deeptools*. Each metaplot represents merged data for the *ITR-1* and *ITR-4* of each accession, except for Hs-0 where only *ITR-1* was detected. **c**, Average size of *Telo*-islands across the *ITR-1* (red) and *ITR-4* (blue) of all accessions, ordered by their total ITR size, as in Extended Data Fig. 7e. As for *CenTel* satellites, the size of *Telo*-islands is more conserved in *ITR-1* than in *ITR-4* domains, possibly due to different emergence and homogenization times or distinct contributions of *ITR-4* HOR types (Col-0, IP-Cat-0 5', and IP-Cat-0 3' types). *Telo*-islands were defined by merging TRs close to each other (<100 bp). **d**, Correlation between ITR size and the number of embedded TRs across accessions. TR number was calculated by identifying and merging all frames of the canonical AAACCCT/TTTAGGG TR motif within the annotated *ITR-1* and *ITR-4* and dividing the length of the resulting sequences by 7. A correlation test (Pearson $R^2$=0.95) between ITR size and TR number shows no significant discrepancies in TR density across ITRs.**e**, Size of large ITRs in the genome assemblies of 110 *A. thaliana* accessions identified using *ITR-Scan*. *ITR-1* and *ITR-4* are displayed in red and blue, respectively. **f**, Sequence similarity dot plot analysis of representative ITR types. ITR sequences from different accessions were compared to themselves and one another using *nucmer* with a 50-bp window, and their sequence similarities were visualized as a heatmap. Comparing each ITR sequence to itself helps visualize the level of similarity between inner ITR segments, defining ITR types, whereas inter-ITR comparisons show the presence or absence of similarity between different ITR types. *ITR-1* type A displays uniform duplications of identical TEs and *CenTel* HORs along the whole ITR, while type B is heterogeneous with several different TEs and *CenTel* HOR organizations. Similar to *ITR-1*, *ITR-4* can be found as type B or type C, a single-HOR ITR usually devoid of TE. While *ITR-1* domains, regardless of their type, show a substantial level of sequence similarity in pairwise comparisons using a 50-bp window, the two *ITR-4* types show a lack of similarity when compared to *ITR-1* domains and to each other. This shows that the two *ITR-4* types differ not only in TE content but also in *CenTel* variants and HOR formation. Several accessions had two *ITR-4*, one of type B and one of type C. **g**, Size of *ITR-1* and *ITR-4* types in the accessions examined. ITR types are depicted in Extended Data Fig. 7e. "B+C" represents the accessions that have two distinct *ITR-4* of types B and C, with each dot representing the summary of the sizes of the two *ITR-4*s. "Complex" corresponds to atypical and highly heterogeneous ITRs.

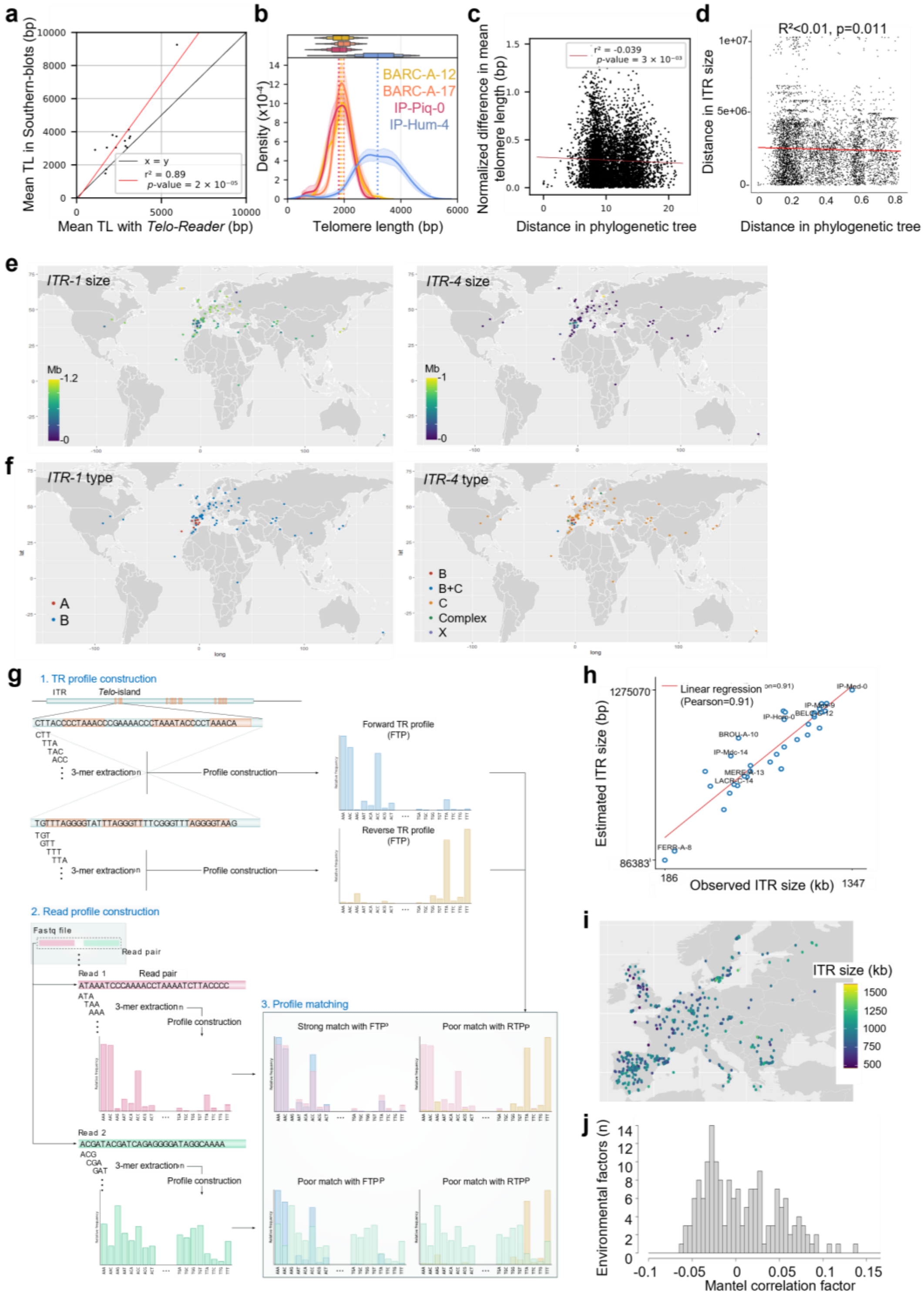


**Extended Data Fig. 8. | Lack of correlation between ITR content and telomere length, phylogeny or environmental variables.**

**a**, Comparison between mean telomere length (TL) distribution computed in this study using long-read sequencing data and previously published Southern blot-based estimates[28], showing a good agreement ($r^2 = 0.89$). **b**, Examples of TL estimates. The TL distributions of two pairs of closely related accessions are shown. BARC-A-12 and BARC-A-17 belong to the same clade and present similar distributions, whereas IP-Piq-0 and IP-Hum-4 also belong to the same clade but present very distinct distributions. The shaded areas around the density curves represent 95% confidence intervals, based on bootstrap sampling of 250 values, with n=100 repetitions with replacement. **c**, Normalized differences in mean TL and phylogenetic distance for all pairs of accessions, showing lack of correlations between the phylogenetic structure and TL across the set of accessions examined. **d**, Pairwise phylogenetic distances in the tree of 110 accessions and differences in ITR length for all pairs of accessions. Correlations were assessed using a Mantel test. The matrices were converted into vectors, and both distances are plotted for each pair of accessions. **e**, Size of *ITR-1* (left) and *ITR-4* (right) across accessions analysed represented on the world map. The majority of the accessions displaying a long *ITR-4* (>100 kb) are found in the Iberian Peninsula. **f**, Architectural types of *ITR-1* (left) and *ITR-4* (right) across all accessions analyzed, represented on the world map. The majority of accessions with type-A *ITR-1* and type-B *ITR-4* are found in the Iberian Peninsula. **g**, Step 1: TR profile construction. 3-mers are extracted from regions over *Telo*-islands within *ITR-1R* and *ITR-4L* of Col-0 and used to construct the forward TR profile, while the reverse complement sequences of the extracted 3-mers were employed to construct a reverse TR profile. Step 2:Read profile construction. Given a FASTQ file, each read pair is processed to construct a profile from the 3-mers extracted from each read in the pair. Step 3: Profile matching. The profiles constructed from each read pair are compared to the forward and reverse TR profiles to determine whether the pair was derived from ITR domains. The figure illustrates an example in which Read 1 matches the forward TR profile. **h**, Benchmarking of ITR length estimation. For 35 accessions with available T2T-assembled genomes, the ITR lengths estimated from NGS data were compared with the corresponding genome coordinates. Since no public NGS data were available for these genomes, we used *art_illumina* simulator to generate artificial paired-end short reads from each assembled genome (read_length=150, fragment_size=250, std=20, cov=40). The artificial sequencing data for each genome were used as input for the k-mer-based method to estimate the ITR lengths. Finally, we calculated the Pearson correlation between the estimated and observed ITR lengths across the 35 genomes. **i**, Detailed view of the geographic distribution of *A. thaliana* accessions of the 1001 Genomes Consortium with their estimated ITR length, complementing Fig. 4f. **j**, Frequency of the correlation factors between the 196 environmental factors given in[50] and ITR lengths of the 531 associated accessions, estimated from the NGS data. Correlations were calculated using a Mantel test correction for phylogenetic distance obtained in this study. None of the factors was significantly related to ITR length. This was then tested using the Pearson correlation factor (without correction for phylogenetic distances) and similar results were obtained ($r<0.15$). Correlation between ITR length and phylogenetic distance was then calculated (without environment) using a Mantel test, and no correlation was observed (Mantel test correlation factor $r= -0.02447588$).

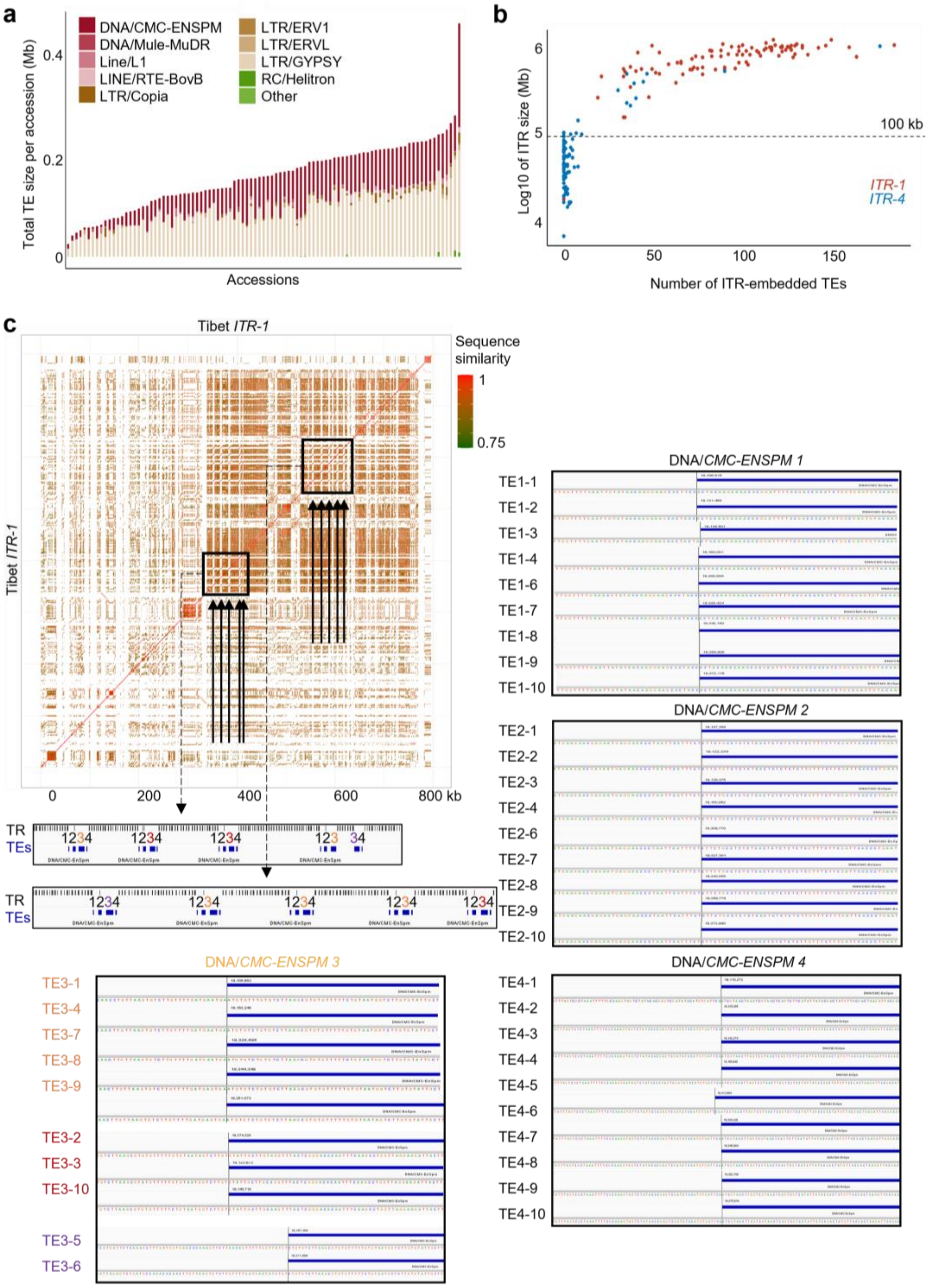


**Extended Data Fig. 9. | The library of ITR-embedded TEs across accessions, and examples in Tibet accession.**

**a**, Representativity of each TE family in the ITRs of each accession plotted as the total size of the annotated TEs belonging to a given TE family. The plot shows the conserved predominance of DNA/*CMC-ENSPM* and LTR/*GYPSY* TEs. **b**, TE number in *ITR-1* (red) and *ITR-4* (blue) for each accession compared to the size of their host ITR (log10 of the length). As illustrated by the dashed bar, ITRs shorter than 100 kb tend to contain few or no TEs. **c**, TE-HOR segment duplicated 9 times along *ITR-1* (black arrows). The duplicated segment comprises four DNA/*CMC-ENSPM* TEs with identical or near-identical integration sites across the 9 loci.

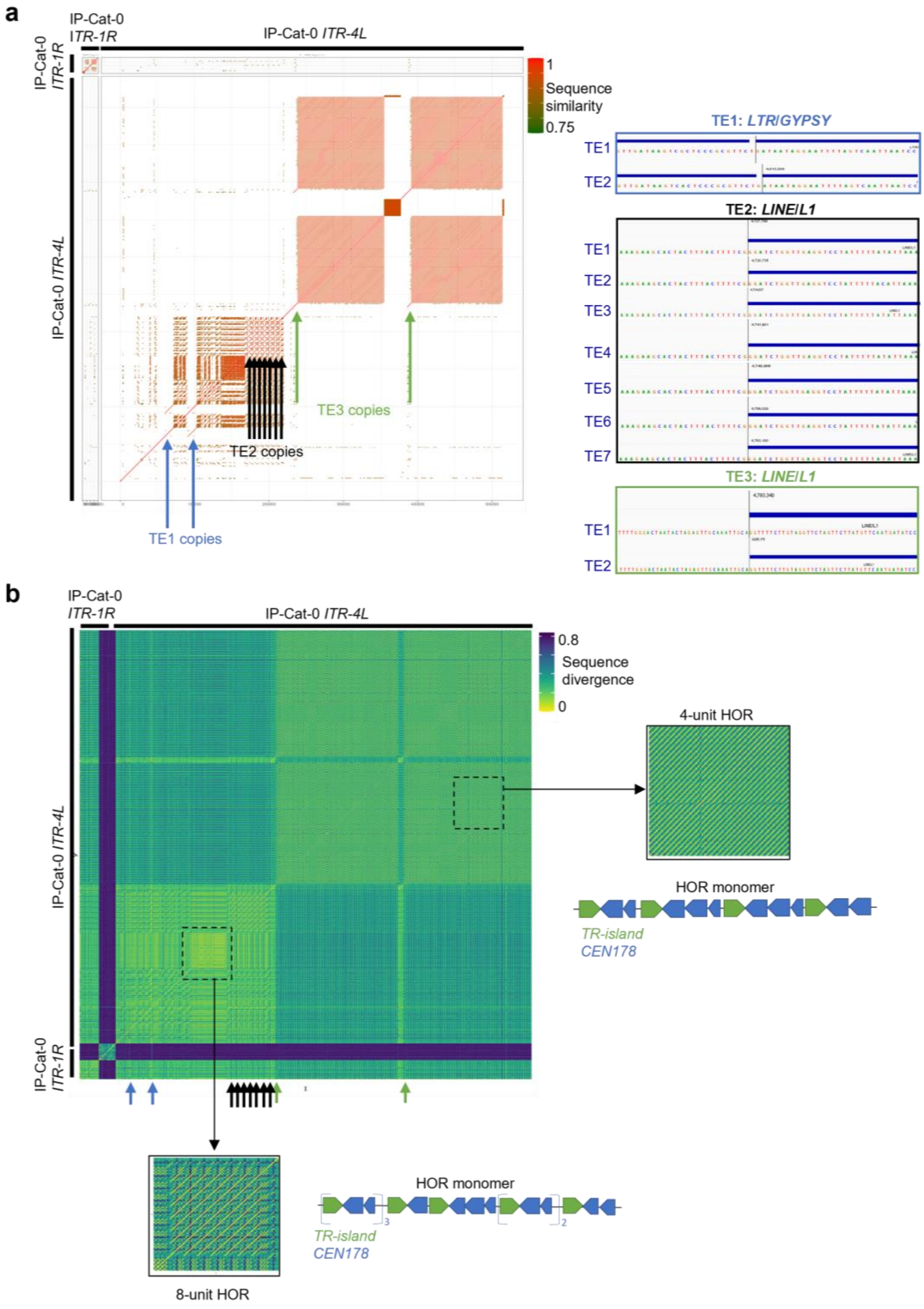


**Extended Data Fig. 10. | TE-associated HOR duplications in IP-Cat-0 accession.**
**a**, Three duplicated TE-HOR segments are shown on a sequence similarity dot plot of *ITR-4*. Duplicated TEs have identical integration sites in all four ITR segments. **b**, Sequence similarity matrix of IP-Cat-0 *CenTel* units, calculated using *ClustalOmega*. Each unit is represented by one matrix cell. Order of units on both axes follows their ITR coordinates. HORs are identified as arrays of *CenTel* satellites with high sequence similarity (yellow diagonal lines). Supporting the joint duplication of TEs with their neighboring ITR segments, duplicated TEs are accompanied by a specific *CenTel* HOR type. To illustrate the methodology, two distinct HORs are augmented, and the *CenTel* units that form the HOR monomers are represented schematically. Color-coded arrows point to TE copies with identical integration sequences.

### Supplementary Tables

**Supplementary Table 1. | Genome sequences analyzed in this study.**

| Description | Sequencing technique | Source |
|---|---|---|
| *TAIR10* | BAC, Sanger | TAIR (GenBank: CP002684.1) |
| *Col-CEN* genome of Col-0 | ONT and PacBio HiFi | Ref. [2] |
| *Col-PEK* genome of Col-0 | ONT and PacBio HiFi | Ref. [51] |
| *Col-XJTU* genome of Col-0 | ONT and PacBio HiFi | Ref. [52] |
| *Col-CC* genome of Col-0 | ONT and PacBio HiFi | *Arabidopsis thaliana* Reference Genome Assembly Consortium (GenBank: GCA_028009825.2) |
| G32 MA lines | PacBio HiFi | This study (ENA Project PRJEB77512) |
| 112 genome assemblies of *A. thaliana* accessions | ONT, PacBio HiFi | Refs [2,35,40–42] |

**Supplementary Table 2. | ChIP-seq, BS-seq, MNase-seq and Hi-C sequencing datasets.**

| Mark | Source | Accession number |
|---|---|---|
| TRB1 | [53] | 10.1038/s41467-023-37263-9 |
| TRB2 | [53] | 10.1038/s41467-023-37263-9 |
| TRB3 | [53] | 10.1038/s41467-023-37263-9 |
| BS-seq (CG, CHG and CHH methyl.) | [54] | 10.1038/s41477-020-00810-z |
| H1.1, H1.2 | [9] | 10.1016/j.celrep.2023.112894 |
| CENH3 | This study | GEO NCBI GSE317061 |
| H3 | [9] | 10.1016/j.celrep.2023.112894 |
| H3K9me2 | [55] | 10.1038/s41467-021-22993-5 |
| H3K27me3 | [9] | 10.1016/j.celrep.2023.112894 |
| H3K4me3 | [56] | 10.1186/s13059-022-02768-x |
| CENH3-OX | [24] | 10.1038/s41586-024-08319-7 |
| Hi-C | [9] | 10.1016/j.celrep.2023.112894 |
| MNase-seq | [58] | GEO NCBI GSE96994 |
| MNase-seq | [59] | GEO NCBI GSE139465 |
| MNase-seq | [60] | GEO NCBI GSE207391 |
| MNase-seq | [61] | GEO NCBI GSE190317 |

**Supplementary Table 3. | Resource data links.**

| Program | Link |
|---|---|
| Telo-Reader | https://github.com/Telomere-Genome-Stability/TeloReader |
| ITR-Scan | https://github.com/i-biocanin/ITRscan |
| ChIP-seq analysis | https://github.com/vidal-adrien/ChIP-Rx-Pipeline-Pub |
| FastaRegexFinder | https://github.com/dariober/bioinformatics-cafe/blob/master/fastaRegexFinder.py |
| Bedtools | https://github.com/arq5x/bedtools2 |
| Bismark v0.24.0 | https://github.com/FelixKrueger/Bismark/releases |
| BLAST | https://blast.ncbi.nlm.nih.gov/Blast.cgi |
| Clustal Omega | https://www.ebi.ac.uk/jdispatcher/msa/clustalo?stype=dna |
| Deepvariant | https://github.com/google/deepvariant |

| | |
|---|---|
| MUMer | https://mummer.sourceforge.net/ |
| Minimap2 | https://github.com/lh3/minimap2 |
| Numpy | https://numpy.org/ |
| Deeptools | https://deeptools.readthedocs.io/en/develop/index.html |
| Picard v2.18 | https://broadinstitute.github.io/picard/ |
| RepeatMasker | https://www.repeatmasker.org/ |
| TRASH | https://github.com/vlothec/TRASH |
| Trimmomatic | http://www.usadellab.org/cms/?page=trimmomatic |
| SciPy | https://scipy.org/ |
| STAR | https://github.com/alexdobin/STAR |
| Samtools | https://github.com/samtools/samtools |
| Sambamba | https://lomereiter.github.io/sambamba/ |
| REPET | https://urgi.versailles.inrae.fr/Tools/REPET/ |
| *ATREP18* annotations | https://www.arabidopsis.org/transposonfamily?key=215 |
| European Nucleotide Archive | https://www.ebi.ac.uk/ena/browser/home |
| fastp v0.22 | https://github.com/OpenGene/fastp |

**Supplementary Table 4. | Detailed analysis and genome coordinates of large ITRs and TEs in 112 *A. thaliana* genome assemblies.** Separate .csv file. Will be given on demand.

**Supplementary Table 5. | ITR genome content estimated from NGS data of 788 *A. thaliana* accessions.** Separate .csv file. Will be given on demand.

**Supplementary Video 1** available at: https://www.pnds-lab.eu/lab-ressources/accessions-itrs
